# In search of Universal Cortical Power Changes Linked to NMDA-Antagonist based Anesthetic Induced Reductions in Consciousness

**DOI:** 10.1101/572636

**Authors:** Andria Pelentritou, Levin Kuhlmann, John Cormack, Steven Mcguigan, Will Woods, Suresh Muthukumaraswamy, David Liley

**Affiliations:** Centre for Human Psychopharmacology, Swinburne University of Technology, Melbourne, Australia; Faculty of Information Technology, Monash University, Melbourne, Australia; Department of Anaesthesia and Pain Management, St. Vincent’s Hospital Melbourne, Melbourne, Australia; Centre for Mental Health, Swinburne University of Technology, Melbourne, Australia; School of Pharmacy, The University of Auckland, New Zealand; Department of Medicine, The University of Melbourne, Parkville, Melbourne, Australia

## Abstract

**Background.:** Despite their intriguing nature, investigations of the neurophysiology of N-methyl-D-aspartate (NMDA)-antagonists Xenon (Xe) and nitrous oxide (N_2_O) are limited and have revealed inconsistent frequency-dependent alterations, in spectral power and functional connectivity. Discrepancies are likely due to using low resolution electroencephalography restricted to sensor level changes, concomitant anesthetic agent administration and dosage. Our intention was to describe the effects of equivalent stepwise levels of Xe and N_2_O administration on oscillatory source power using a crossover design, to explore universal mechanisms of NMDA-based anesthesia.

**Methods.:** 22 healthy males participated in a study of simultaneous magnetoencephalography and electroencephalography recordings. In separate sessions, equivalent subanesthetic doses of gaseous anesthetic agents N_2_O and Xe (0.25, 0.50, 0.75 equi MAC-awake) and 1.30 MAC-awake Xe (for Loss of Responsiveness) were administered. Source power in various frequency bands was computed and statistically assessed relative to a conscious baseline.

**Results.:** Delta (l-4Hz) and theta (4-8Hz) band power was significantly increased at the highest Xe concentration (42%, 1.30 MAC-awake) relative to baseline for both magnetoencephalography and electroencephalography source power (p<0.005). A reduction in frontal alpha (8-13 Hz) power was observed upon N_2_O administration, and shown to be stronger than equivalent Xe dosage reductions (p=0.005). Higher frequency activity increases were observed in magnetoencephalographic but not encephalographic signals for N_2_O alone with occipital low gamma (30-49Hz) and widespread high gamma (51-99Hz) rise in source power.

**Conclusions.:** Magnetoencephalography source imaging revealed unequivocal and widespread power changes in dissociative anesthesia, which were divergent to source electroencephalography. Loss of Responsiveness anesthesia at 42% Xe (1.30 MAC-awake) demonstrated, similar to inductive agents, low frequency power increases in frontal delta and global theta. N_2_O sedation yielded a rise in high frequency power in the gamma range which was primarily occipital for lower gamma bandwidth (30-49 Hz) and substantially decreased alpha power, particularly in frontal regions.

**Clinical trial number and Registry URL:** Not applicable.

**Prior Presentations:** Pelentritou Andria, Kuhlmann Levin; Lee Heonsoo; Cormack John; Mcguigan Steven; Woods Will; Sleigh Jamie; Lee UnCheol; Muthukumaraswamy Suresh; Liley David. Searching For Universal Cortical Power Changes Linked To Anesthetic Induced Reductions In Consciousness. The Science of Consciousness April 4^th^ 2018. Tucson, Arizona, USA.

**Summary Statement:** Not applicable.

## 1. Introduction

Though effectively and extensively used in medical interventions on a daily basis, the molecular, neural and behavioral changes underlying anesthetic induced unconsciousness remain elusive. This is less true of general anesthetics that are primarily gamma-aminobutyric acid (GABA) receptor agonists like propofol and volatile agents^1,2^. The dissociative agents ketamine, N_2_O and Xe have a different pharmacological profile, being predominately NMDA receptor antagonists with weak effects on the GABA receptor^3-6^. The distinction between the two classes of drugs extends to their electrophysiology which are relevant for Depth of Anesthesia monitors. Electroencephalography under propofol anesthesia is well characterized and reveals global changes in brain activity associated with the appearance of high-amplitude, low-frequency delta waves, frontal alpha oscillations and a persistent reduction in higher frequencies, notably gamma activity^7-10^. In contrast, the electroencephalographic effects of NMDA based agents which tend to yield variable effects with ketamine and N_2_O generally increasing high frequency gamma power^11-14^ and having no effects or decreasing low frequency power^13,14^. Conflicting evidence suggests that N_2_O sustains gamma and alpha activity and instead increases delta and theta power^15,16^. Such ambiguity extends to Xe anesthesia wherein topologically diverse increases and decreases in delta^17^, theta^17,18^, alpha^18,19^ and gamma^17,20^ activity have been reported.

The anomalous electrophysiology of the NMDA antagonistic anesthetics may be attributed to a number of factors. Firstly, investigations of gaseous N_2_O and Xe when compared to intravenous ketamine, are limited to a small number of studies potentially due to the unclear clinical utility of Xe as well as the widely reported association with post-operative nausea and vomiting^21-23^, and pronounced dissociative effects. In addition, dosage differences across studies ranging from subanesthetic to anesthetic levels^14,15,19,20,24^ and the common practice inclusion of concomitant agents^17,25^, encumber the ability to draw conclusive findings. Acquiring data has also been limited to electroencephalography recordings from depth of anesthesia monitors or other spatially low resolution systems^12,26,27^, which makes the investigation of network level changes and mesoscopic processes difficult. While electroencephalography is extremely useful, it suffers from the problem of volume conduction and limited spatial resolution even in high density systems, all of which can be better solved by implementing source level electroencephalography and magnetoencephalography imaging^28^. When used in conjunction, the two techniques are highly complementary, recording from both radial and tangential sources in the brain and picking up a wider range of activity, particularly in the high-frequency range^29^. However, only a handful of studies have utilized the method to study anesthetic effects^30-32^ and have mostly focused on task-related changes^30,31^.

The idiosyncrasies underlying the action of the gaseous NMDA antagonists Xe and N_2_O with the former seemingly resembling the GABA agonists in electrophysiology, may enhance the study of anesthetic induced unconsciousness and offer novel insights into associated neural correlates. Therefore, this study measured the electromagnetic effects of subanesthetic administration of Xe and N_2_O in healthy volunteers using source level brain imaging and spectral analysis, in attempting to outline similarities and differences between the two agents, contrasted to wakefulness.

## 2. Materials and Methods

The study entitled “Effects of inhaled Xe and N_2_O on brain activity recorded using EEG and MEG” was approved (approval number: 260/12) by the Alfred Hospital and Swinburne University of Technology Ethics Committee and met the requirements of the Australian National Statement on Ethical Conduct in Human Research (2007).

### Study Population

Twenty-two volunteers were recruited from November 2015 to November 2017 using recruitment flyers posted throughout Swinburne University of Technology. Phone screening was conducted by a member of the research team to ensure eligibility for participation. One volunteer was excluded following participation due to a conflicting medical condition revealed to the neuroimaging facility neurologist upon detailed review of the relevant MRI scan. All participants signed a written informed consent and were given monetary compensation for their time (300 AUD).

Participants were regarded as eligible if they were right handed, adult males, between the ages of 20 and 40 (mean age: 24), had a Body Mass Index (BMI) between 18-30 and were American Society of Anesthesiologists (ASA) Physical status 1 in accordance with the day stay general anesthesia procedure as designated in the Australia and New Zealand College of Anaesthetists (ANZCA) guidelines (Document PS15). Females were excluded due to the documented effects of menstruation^33^ and/or age extremes on the resting MEG/EEG signal as well as the increased propensity to nausea and vomiting^34^. Contraindications to MRI or MEG (such as implanted metallic foreign bodies) were an additional exclusion criterion. Exclusion from the study resulted if candidates had neurological disorders, psychiatric disorders, epilepsy, heart conditions, respiratory conditions, obstructive sleep apnea, asthma, motion sickness and claustrophobia. In addition, any recent intake of psychoactive or other prescribed medication as well as any recreational drug use resulted in non-inclusion. An inventory of previous surgeries was taken upon screening, importantly any unfavorable reactions to general anesthesia which would result in exclusion from the study. Finally, to ensure a good seal with an anesthetic face mask, participants with large beards were excluded unless they were willing to shave.

### Anesthetic Protocol

A detailed description of the data acquisition protocol with accompanying video documentary has been published previously^35^. Here, the key components are summarized.

A two-way crossover experimental design was followed. Two separate testing sessions were performed for each subject separated by a maximum of four weeks, with Xe being administered in one session and N_2_O in a second session. Participants were blind to the type of gas being administered while the medical staff and researchers were not due to the subtle differences in the procedure followed for their administration and to safely administer the study gas. At least one anesthesiologist was present in the control room to manage gas delivery and electronic monitoring and an anesthetic nurse or other suitably trained clinical observer sat with the subject in the magnetically shielded room containing the magnetoencephalography and electroencephalography equipment to effectively monitor the participant’s condition (in particular the face mask seal and subject’s airway patency).

In line with the day stay ANZCA guidelines (document PS15), volunteers were asked to fast for at least six hours and consume no liquids for at least two hours prior to the start of the experiment. Participant eligibility was reaffirmed with an extensive medical history interview and vital sign measurements which include blood pressure, heart rate, body temperature and peak expiratory flow followed by a brief measurement in the MEG to ensure that there are no unanticipated sources of noise. A peripheral intravenous catheter was placed to enable anti-emetic administration consisting of 4mg dexamethasone and 4mg ondansetron to reduce the incidence of emesis caused by anesthetic gas inhalation^22^, which is often observed with N_2_O at the higher concentrations used^21,36^. A face mask and breathing circuit was attached to the subject using a modified sleep apnea continuous positive airways pressure (CPAP) harness. Anesthesia gases were delivered using the Akzent Xe Color anesthesia machine (Stephan GmbH, Gackenbach, Germany), located outside the magnetically shielded room with anesthesia hoses passing through magnetically shielded room portals. End-tidal Xe concentrations are measured using katharometry and N_2_O concentrations are measured using infrared spectroscopy implemented in the anesthesia machine. Prior to anesthetic administration, each participant inhaled 100% inspired O_2_ for up to 30 minutes until their end-tidal O_2_ concentration was >90%, effectively de-nitrogenating the participant in order to ensure effective end-tidal anesthetic gas concentrations.

In addition to magnetoencephalography and electroencephalography recordings (see Data Acquisition), three bipolar bio-channel recordings were made: electromyography to record the activity of the mylohyoid and digastric (anterior belly) muscles, electrooculography and three-lead electrocardiography. Pulse oximetry and non-invasive blood pressure measurement (NIBP) was registered outside the magnetically shielded room as per ANZCA Guidelines (Document PS18). The ongoing level of responsiveness was behaviorally measured throughout the experiment using an auditory continuous performance task in which a binaural auditory tone of either 1 or 3 kHz frequency of fixed stereo amplitude (approximately 76 dB), with an inter-stimulus interval of between 2 to 4 seconds drawn from a uniform distribution. Participants responded using two separate button boxes held in each hand. The left and right buttons on each box correspond to a low or high frequency tone, respectively, and the left and right button boxes were respectively used for the participant to indicate the absence or presence of nausea. Three successive button presses indicating nausea or signs of nausea as identified by the anesthesiologist or clinical observer were evaluated and if severe, gas inhalation was terminated.

All participants underwent the following recordings with magnetoencepholagram and electroencephalogram data collection throughout each period. Three eyes closed baseline conditions, all prior to gas administration: A 5-minute resting period (baseline 1) followed by a 5-minute period with the auditory continuous performance task being performed (baseline 2) and a final 5-minute recording following anti-emetic administration with the auditory continuous performance task being performed (baseline 3). Four step-wise increasing levels for Xe administration and three step-wise increasing levels for N_2_O administration: MAC-awake levels of 0.25 (level 1), 0.5 (level 2) and 0.75 (level 3) times the MAC-awake concentration with concentrations at 8 %, 16 %, 24 % and 1 6%, 32 %, 47 % concentrations for Xe/ O_2_ and N_2_O / O_2_, respectively. At each level, a 10 minute gas wash-in took place such that the target end-tidal gas concentration was reached, a 5 minute steady-state period ensued followed by a final 10 minute wash-out period with the administration of 100 % O_2_ during which end-tidal gas concentration returns to 0. A fourth level was considered for Xe corresponding to 1.3 (level 4) times the MAC-awake concentration (42 % Xe/O_2_) upon which 95 % of participants are expected to lose responsiveness. In our experiment, 76 % of volunteers reached the loss of responsiveness state (see results for details on what happened to further excluded participants). A fourth level was not included for N_2_O administration as it is known that high levels induce nausea and vomiting, and are associated with greater risk of hypoxia at high concentrations. Once Loss of Responsiveness was achieved at Xe level 4, the Xe gas level was maintained for up to 10 minutes or until the medical staff considered airway patency was compromised, after which wash-out with 100 % O_2_ took place.

Participants were discharged once they were awake, alert, and responsive, without significant nausea or vomiting, were able to ambulate with minimal assistance, and had a responsible adult to accompany them home. They were asked to complete a truncated version of the 5-Dimensional Altered States of Consciousness Rating Scale (5D-ASC) questionnaire^37,38^ and a short narrative of their overall experience during the experiment as well as specific details about level dependent qualitative effects, both of which were performed 24 hours after each recording session. Cognitive and subjective report data collected are not discussed here.

### Data Acquisition

#### Electrophysiological Recordings

Data acquisition was simultaneously performed in a room shielded from magnetic or electric interferences (Euroshield Ltd., Eura, Finland). Brain magnetic field activity (MEG) was recorded at a sampling rate of 1000 Hz using a Helmet-shaped TRIUX Whole-head 306-channel magnetometer system (Elekta Neuromag306, Oy, Helsinki, Finland) consisting of 204 planar gradiometers and 102 magnetometers. Head position in relation to the recording system was recorded using 5 head-position indicator coils and was continuously monitored by measuring the magnetic fields produced by the coils in relation to the cardinal points on the head (nasion, left and right pre-auricular points) which were determined prior to commencement of the experiment using Isotrack 3D digitizer (Polhemus, Colchester, VT, USA). The internal active shielding system for threedimensional noise cancellation was disabled to allow for subsequent source space analysis.

Electroencephalogram data were acquired at a sampling rate of 512 Hz using an MEG compatible 64-channel Waveguard cap (ANT Neuro, Enschede, Netherlands), an ASALAB battery powered amplifier (ANT Neuro, Enschede, Netherlands), a magnetically shielded cordless battery box and ASALAB acquisition software (ANT Neuro, Enschede, Netherlands). Electrical contact impedances were kept below 5 kO.

#### Structural Tl-Magnetic Resonance Imaging

A single structural T1-weighted Magnetic Resonance Imaging scan was performed using the 3.0 TIM Trio MRI system (Siemens AB, Erlangen, Germany) with markers used to highlight the digitized fiducial points for the nasal apex and left and right pre-auricular points for subsequent source space reconstruction. T1-weighted images were acquired on a sagittal plane with a magnetization prepared rapid gradient echo (MP-RAGE) pulse sequence with an inversion recovery (176 slices, slice thickness = 1.0 mm, voxel resolution = 1.0 mm^3^, TR = 1900 ms, TE = 2.52 ms, TI = 900 ms, bandwidth = 170 Hz/Px, flip angle = 9°, field of view 350 mm x 263 mm x 350 mm, orientation sagittal, acquisition time = ~ 5 min).

### Data Analysis

#### Preprocessing

Data analysis was performed in Fieldtrip version 20170801^39^ and custom MATLAB (MathWorks, Natick, MA, USA) scripts and toolboxes. MEG data from 21 subjects and EEG data from 19 subjects (one dataset missing due to an administrative error and a second excluded due to issues with forward model construction, potentially due to a corrupt electroencephalography digitization file) were visually inspected to exclude any malfunctioning channels from further analysis. MEG data was filtered using the temporal signal-space separation algorithm^40^ of Max Filter software version 2.2 (Elekta Neuromag Oy, Helsinki, Finland) and the signal from magnetometers and planar gradiometers combined in Fieldtrip using provided algorithms. The two datasets were time adjusted for any temporal differences in acquisition using a synchronization trigger. Given the EEG and MEG machines did not share the exact same electronic clock, meaning that synchronization of the two data types had a small margin of error, the EEG and MEG data were analyzed separately. The relevant periods of interest were selected for EEG and MEG data which included the three 5 minute baselines, three 5 minute anesthetic steady-state periods and the entire loss of responsiveness period for Xe which were then band-pass filtered at 1 to 100 Hz and any line noise at 50, 100 and 150 Hz removed. EEG data were re-referenced using the common average reference method. All periods of interest were visually inspected and any artefactual segments resulting from eye movements or blinks, jaw clenches or movements, head movements, breathing and other muscle artefacts were excluded from further analysis. Six classical bandpass-filtered versions of the datasets were created for subsequent source analysis: delta (1-4Hz), theta (4-8Hz), alpha (8-13Hz), beta (13-30Hz), low gamma (30-49Hz) and high gamma (51-99Hz).

#### Source localization

Co-registration of each subject’s MRI to their scalp surface was performed using fiducial realignment and boundary element method volume conduction models were computed using a single shell for the MEG^41^ and best fit dipole orientation for the EEG^42^. Both were spatially normalized to a template MRI using the Segment toolbox in SPM8^43^to serve as a volume conduction model in subsequent source level analyses. An atlas-based version^44^ of the Linearly Constrained Minimum Variance (LCMV) spatial filtering beamformer^45^ was used to project sensor level changes onto sources. Global covariance matrices for each band (after artefact removal) were derived which were utilized to compute a set of beamformer weights for all brain voxels at 6mm voxel resolution. Following inverse transformation, the 5061 voxels were assigned one of ninety AAL atlas labels (78 cortical, 12 subcortical)^46^ in the subjects’ normalized co-registered MRI (based on proximity in Euclidean distance to centroids for each region) in order to reveal regional anesthetic-induced changes in oscillatory power. The resulting beamformer weights were normalized to compensate for the inherent depth bias of the method used^44^. Finally, the original time-series were segmented into contiguous 3 second epochs (approximately in accordance with epochs used in other studies^47^) and the final epoch covariance matrices were used along with normalized weights to compute the power at each voxel across all frequencies within a band^45^.

### Statistical Analysis

In order to ensure equal numbers of epochs across subjects, gases and gas levels, epochs from larger datasets were selected at random, resulting in a total of 17 three second epochs (51 seconds - 51000 samples) for magnetoencephalography data and a total of 9 three second epochs (27 seconds - 13824 samples) for electroencephalography data being used for non-parametric group statistical analysis within each frequency band. The discrepancy in final epoch numbers was due to the independent artefact removal performed on the magnetoencephalography and electroencephalography recordings, the latter being more sensitive to eye blinks and jaw movements. In addition, the variation in filtering techniques could influence the presence of artefactual segments with max-filtering (as described earlier) applied to the magnetoencephalogram being a powerful and highly effective additional filter not applicable for electroencephalography data^40^.

#### Within Gas Comparison

For each individual, student’s t-statistic images of the source power at every voxel were calculated for the two pre-antiemetic baselines and all gas levels versus the post-antiemetic baseline, to allow for comparative imaging. In order to reject the null hypothesis stating that a given condition in either the Xe or N_2_O case is not significantly different to the post-antiemetic baseline, a null distribution was constructed using five thousand permutations of the t-statistic sign and a one sample t-test was finally used to compute the effect at each voxel across individuals^48,49^. Maximum statistics with significance level p=0.05 were utilized to correct for the voxel multiple comparisons problem^48^ and Bonferroni correction was further considered to compensate for multiple comparisons between each condition and the eyes-closed post-antiemetic baseline. This resulted in a threshold of p=0.004 for the Xe conditions and p=0.005 for the N_2_O conditions (p=0.025 to allow for two tailed comparison of increases and decreases in power; p=0.025/6 and p=0.025/5 to correct for the six Xe and five N_2_O condition comparisons, respectively).

#### Across Gas Comparison

To compare the effects of the two anesthetic agents, student’s t-statistic images were computed for each of three baselines and three equi MAC-awake conditions of N_2_O against Xe on each individual’s source power. In order to reject the null hypothesis stating that there is no significant difference in a given condition between Xe or N_2_O, a null distribution was constructed using five thousand permutations of the t-statistic sign and a one sample t-test was finally used to compute the effect at each voxel across individuals^48,49^. Maximum statistics with p=0.05 were utilized to correct for the voxel multiple comparisons problem^48^ and Bonferroni correction was used to compensate for the multiple comparisons of considering each condition (3 baselines and 3 levels). This resulted in a threshold of p=0.005 for the Xe and N_2_O conditions (p=0.025 to allow for two tailed comparison of increases and decreases in power; p=0.025/6 and p=0.025/6 to correct for the six Xe and N_2_O condition comparisons).

Figure 1 offers an overall view of the experimental design and briefly summarizes both data acquisition and data analysis for this study.

**Figure 1.**
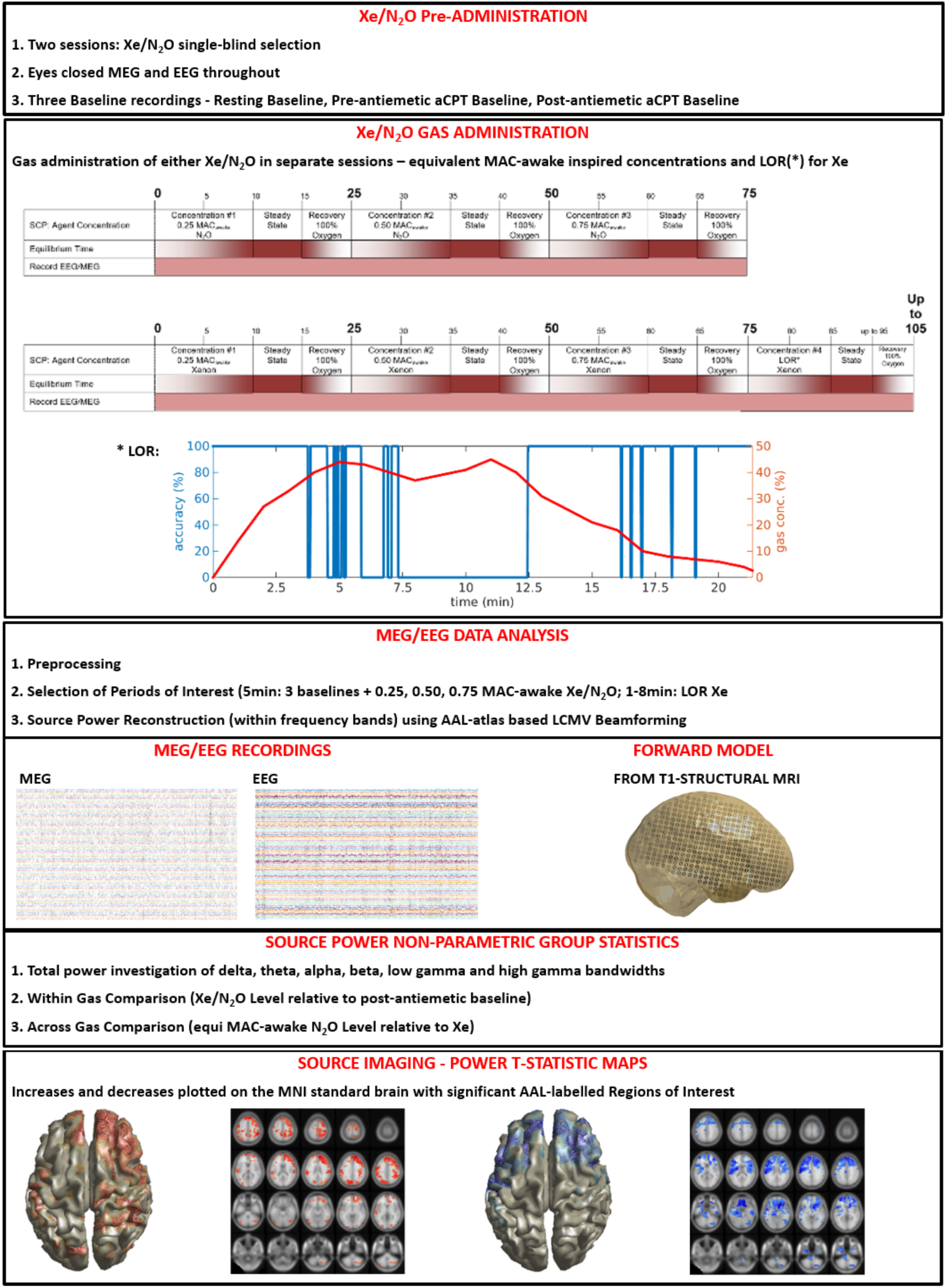
Study Synopsis. Illustration of the data acquisition, magnetoencephalography (MEG) and electroencephalography (EEG) data analysis and statistical analysis. [aCPT: auditory continuous performance task; LOR: Loss of Responsiveness; delta (1-4 Hz), theta (4-8 Hz), alpha (8-13 Hz), beta (13-30 Hz), low gamma (30-49 Hz), high gamma (51-99 Hz)].

### 3. Results

#### Anesthetic Protocol Outcomes

All twenty-one participants included in the analysis successfully completed the 0.25, 0.50, 0.75 MAC-awake N_2_O sub-anesthetic levels and completed the 0.25, 0.50 MAC-awake Xe sub-anesthetic level. One participant was excluded from further Xe analysis as he felt unable to continue beyond 0.50 MAC-awake Xe therefore, twenty participants accomplished 0.75 MAC-awaked Xe anesthesia. Of the remaining twenty, sixteen participants achieved loss of responsiveness (LOR) at the final 1.30 MAC-awake Xe concentration while the remaining four failed to meet this criterion at the highest level due to gas leakage or excessive nausea. The period of loss of responsiveness varied across the sixteen individuals between 60 and 480 seconds (mean=213.25 seconds). All remaining periods of interest for baselines and gas levels were 300 seconds. All MEG datasets were utilized while two EEG datasets were excluded from the analysis one. Please refer to table 1A in Supplementary Digital Content 1 which summarizes the above for clarity.

**Table 1.** Magnetoencephalographic sources most significantly altered byXe and N_2_0 administration. A selection of five representative regions of interest was picked from the twenty highest t-value and p-value (p<0.005 for N_2_O; p<0.004 for Xe) regions for each frequency band. Region of interest analysis reveals most pronounced increases for delta in frontal regions and a widespread theta rise relative to baseline during Xe Loss of Responsiveness (1.30 MAC-awake). The highest N_2_O inspired concentration (0.75 MAC-awake) resulted in primarily frontal alpha reductions and effects on low gamma were primarily observed in occipital voxels while high gamma was localized to a small number of frontal and occipital regions. Voxel coordinates are in AAL atlas coordinate system along with associated labels^46^, [delta (1-4 Hz), theta (4-8 Hz), alpha (8-13 Hz), low gamma (30-49 Hz), high gamma (51-99 Hz)].

| Frequency Band | MAC-awake Level | Region of Interest | Voxel Coordinate | t-value | p-value |
| --- | --- | --- | --- | --- | --- |
| **delta** | 1.30 Xe | Parietal_Inf_R | 54 -42 48 | 9.61 | 0.0001 |
|  | 1.30 Xe | Frontal_Sup_Medial_L | 0 54 24 | 9.25 | 0.0002 |
|  | 1.30 Xe | Frontal_Sup_Medial_R | 12 48 24 | 9.13 | 0.0002 |
|  | 1.30 Xe | Precuneus_L | 0 -48 54 | 9.13 | 0.0002 |
|  | 1.30 Xe | Frontal_Sup_R | 18 54 30 | 9.11 | 0.0002 |
| **theta** | 1.30 Xe | Parietal_Sup_L | -30 -66 48 | 8.52 | 0.0003 |
|  | 1.30 Xe | Precuneus_L | -6 -78 54 | 8.40 | 0.0004 |
|  | 1.30 Xe | Frontal_Sup_R | 24 24 60 | 8.20 | 0.0004 |
|  | 1.30 Xe | Parietal_Sup_R | 18 -60 54 | 8.15 | 0.0004 |
|  | 1.30 Xe | Precuneus_R | 12 -66 42 | 8.09 | 0.0004 |
| **alpha** | 0.75 N<sub>2</sub>O | Frontal_Sup_Medial_L | 0 66 0 | -9.92 | 0.0001 |
|  | 0.75 N<sub>2</sub>O | Frontal_Mid_L | -42 54 6 | -9.70 | 0.0002 |
|  | 0.75 N<sub>2</sub>O | Frontal_Inf_Tri_L | -48 36 18 | -9.60 | 0.0002 |
|  | 0.75 N<sub>2</sub>O | Frontal_Sup_Orb_R | 12 18 -18 | -9.38 | 0.0002 |
|  | 0.75 N<sub>2</sub>O | Frontal_Inf_Orb_R | 30 30 -24 | -8.83 | 0.0002 |
| **low gamma** | 0.75 N<sub>2</sub>O | Occipital_Sup_L | -12 -90 30 | 9.58 | 0.0001 |
|  | 0.75 N<sub>2</sub>O | Occipital_Mid_R | 30 -90 18 | 9.21 | 0.0002 |
|  | 0.75 N<sub>2</sub>O | Occipital_Sup_R | 24 -84 12 | 8.84 | 0.0002 |
|  | 0.75 N<sub>2</sub>O | Occipital_Inf_L | -24 -96 -6 | 8.50 | 0.0002 |
|  | 0.75 N<sub>2</sub>O | Occipital_Mid_L | -30 -84 36 | 8.42 | 0.0002 |
| **high gamma** | 0.75 N<sub>2</sub>O | Frontal_Mid_R | 24 54 30 | 7.18 | 0.0001 |
|  | 0.75 N<sub>2</sub>O | Frontal_Sup_Medial_L | -6 30 60 | 6.94 | 0.0002 |
|  | 0.75 N<sub>2</sub>O | Precuneus_R | 6 -78 48 | 6.92 | 0.0002 |
|  | 0.75 N<sub>2</sub>O | Frontal_Sup_R | 24 54 36 | 6.83 | 0.0002 |
|  | 0.75 N<sub>2</sub>O | Cingulum_Mid_L | 0 0 42 | 6.81 | 0.0002 |

#### Preliminary Spatially Averaged Source Level Total Power Analysis

Total power and relative changes [(Gas power-Baseline power)/Baseline power) x 100] across all subjects and across all voxels was compared to post anti-emetic baseline (baseline 3). This was performed within each gas type, within inspired gas concentration, within each of the six frequency ranges investigated with ±1 Standard Deviation (See Figure 2, error bars) and all results are indicated in Figure 2. Table 1B in Supplementary Digital Content 1 gives the corresponding numerical values and ±1 Standard Deviation (SD) for all differences briefly summarized here. Magnetoencephalography findings demonstrate increases in low frequency delta, theta and beta power at the highest levels of Xe anaesthesia. Reductions in alpha power and increases in high frequency gamma power were observed at the highest inspired doses of both Xe and N_2_O. Electroencephalography power showed increases in both gases at highest dosage of Xe and N_2_O. Decreased electroencephalographic alpha power was again observed but was more prominent at the intermediate 0.50 MAC-awake dosage compared to higher 0.75 MAC-awake. High frequency activity, like magnetoencephalography results, showed increases in the highest Xe and N_2_O inspired concentrations. Preliminary results revealed that the electromagnetic effects of anesthesia depend on agent administered and importantly modality measured and hence, we chose to independently compare the two in subsequent analysis and corresponding results.

**Figure 2.**
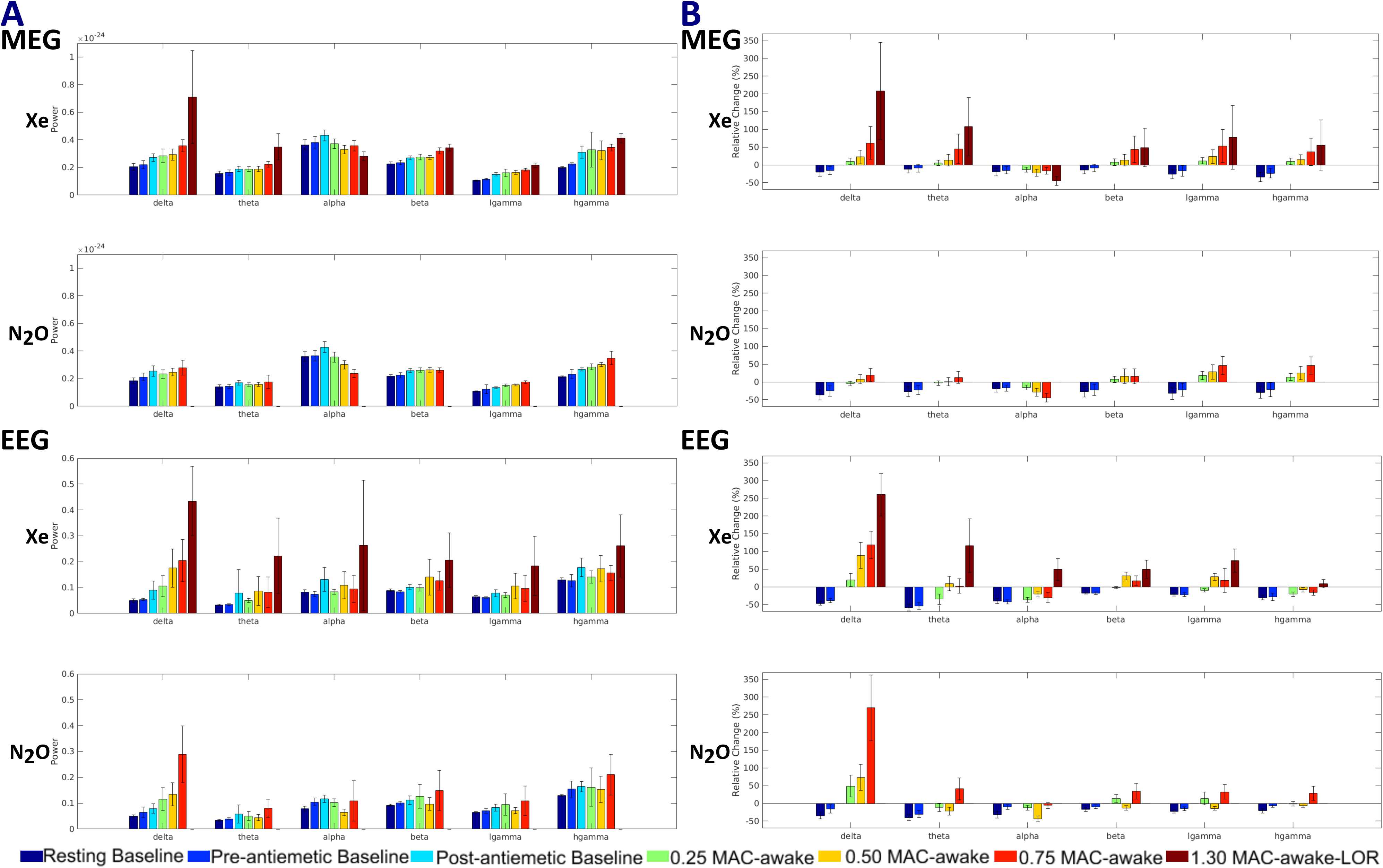
Total power alterations under Xe and N_2_0 anesthesia relative to post-antiemetic baseline in various frequency bands. Baseline and Gas power (A) at each concentration is averaged across all samples for all subjects and considering all voxels of the magnetoencephalogram (MEG) and electroencephalogram (EEG). Units of power for MEG (A) are T^2^ and for EEG (B) are |iV^2^. The figure demonstrates changes in all frequency bands that are different for the two modalities and in some cases correlate to increasing gas concentration. Error bars illustrate ±1 SD. Relative change (B) ([(Gas or Baseline power-Post-antiemetic Baseline power)/Post-antiemetic Baseline power] x 100) is calculated and all changes are shown with each sub-figure in bold percentage values, [delta (1-4 Hz), theta (4-8 Hz), alpha (8-13 Hz), beta (13-30 Hz), low gamma (30-49 Hz), high gamma (51-99 Hz)].

Upon evaluation of magnetoencephalographic power, increases relative to baseline were seen in the delta (relative change: 208%, SD: 136% at Loss of Responsiveness) and theta bands (relative change: 107%, SD: 82% at Loss of Responsiveness) for higher concentrations of Xe. No changes were seen in the delta and theta power for N_2_O. In addition, a minor increase in the beta band was observed at the highest Xe concentration (relative change: 24.9%, SD: 6.71%). Both gases decreased alpha power compared to baseline (relative change: Xe at Loss of Responsiveness: −44.6%, SD: 12.9%; Xe at 0.75 MAC-awake: −16.8%, SD: 9.36%; N_2_O at 0.75 MAC-awake: −44.4%, SD: 12.6%) and increased gamma activity, particularly in the high frequency range (51-99Hz) (relative change: low gamma: Xe at Loss of Responsiveness: 77.5%, SD: 89.9%; N_2_O at 0.75 MAC-awake: 46.3%, SD: 25.6%; high gamma: Xe at Loss of Responsiveness: 55.0%, SD: 71.7%; N_2_O at 0.75 MAC-awake: 45.7%, SD: 24.5%).

Electroencephalography results showed similar trends to those of magnetoencephalography in lower frequency activity however, it has to be noted that total power displayed larger variability throughout (see Supplementary Digital Content 1, Table 1B for further details) particularly in the highest 1.30 MAC-awake Xe concentration and in the lower delta and theta bandwidths. Nonetheless, delta band increases were observed for all stepwise increasing concentrations of both gases relative to baseline (relative change: Xe at Loss of Responsiveness: 260%, SD: 60%; N_2_O at 0.75 MAC-awake: *270%,* SD: 92.5%). Xe power alone was substantially higher at Loss of Responsiveness (1.30 MAC-awake Xe) for the theta band (relative change: 116%, SD: 75.8%). A reduction in alpha power is again observed in electroencephalographic sources after N_2_O administration that appears larger for 0.50 MAC-awake compared to the higher 0.75 MAC-awake concentration (relative change: N_2_O at 0.75 MAC-awake: −5.71%, SD: 9.05% vs. N_2_O at 0.50 MAC-awake: −44.0%, SD: 8.10%). Interestingly, Xe alpha power was increased at Loss of Responsiveness (relative change: Xe at Loss of Responsiveness: 49.3%, SD: 30.9% vs. N_2_O at 0.50 MAC-awake: −44.0%, SD: 8.10%), in contrast to magnetoencephalography relative power changes. This deviation between measurement types and any dissimilarities observed for higher frequency activity likely reflect the high degree of variation in the data.

### Within Gas Statistical Analysis

Power changes were evaluated at each gas level relative to the post-antiemetic baseline (baseline3). Since our focus was on reliable statistically significant changes, we controlled significance by combined corrections for multiple comparisons across space using maximum statistics and across conditions using Bonferroni correction. It should be noted that significant maximum statistics corrected (without subsequent Bonferroni correction) t-statistic maps of the power changes relative to baseline across subjects displayed potential trends in the data with increasing gas concentration (see Supplementary digital content 2 and Figure 2 for magnetoencephalography and electroencephalography recordings). As revealed in the preliminary analysis, Xe and N_2_O induced power changes depended not only on the anesthetic agent used but also on the electrophysiological modality. While magnetoencephalography source power changes spanned the whole frequency spectrum being investigated, electroencephalography alterations were prominent only in lower frequencies.

#### Magnetoencephalography Results

Figure 3 encapsulates the most significant power changes in electromagnetic activity and table 1 displays a selection of corresponding regions of interest (see Supplementary Digital Content 2, Table 2A for complete list of significant regions of interest). Overall, Xe effects were specific to low delta and theta band frequencies with no changes observed on high frequency power. Conversely, N_2_O did not significantly alter low frequency delta and theta activity but rather increased high frequency low and high gamma activity. Effects on the alpha band were specific to N_2_O administration which induced reductions in alpha power, most prominent in frontal regions. Neither anesthetic significantly altered beta power.

**Table 2.**
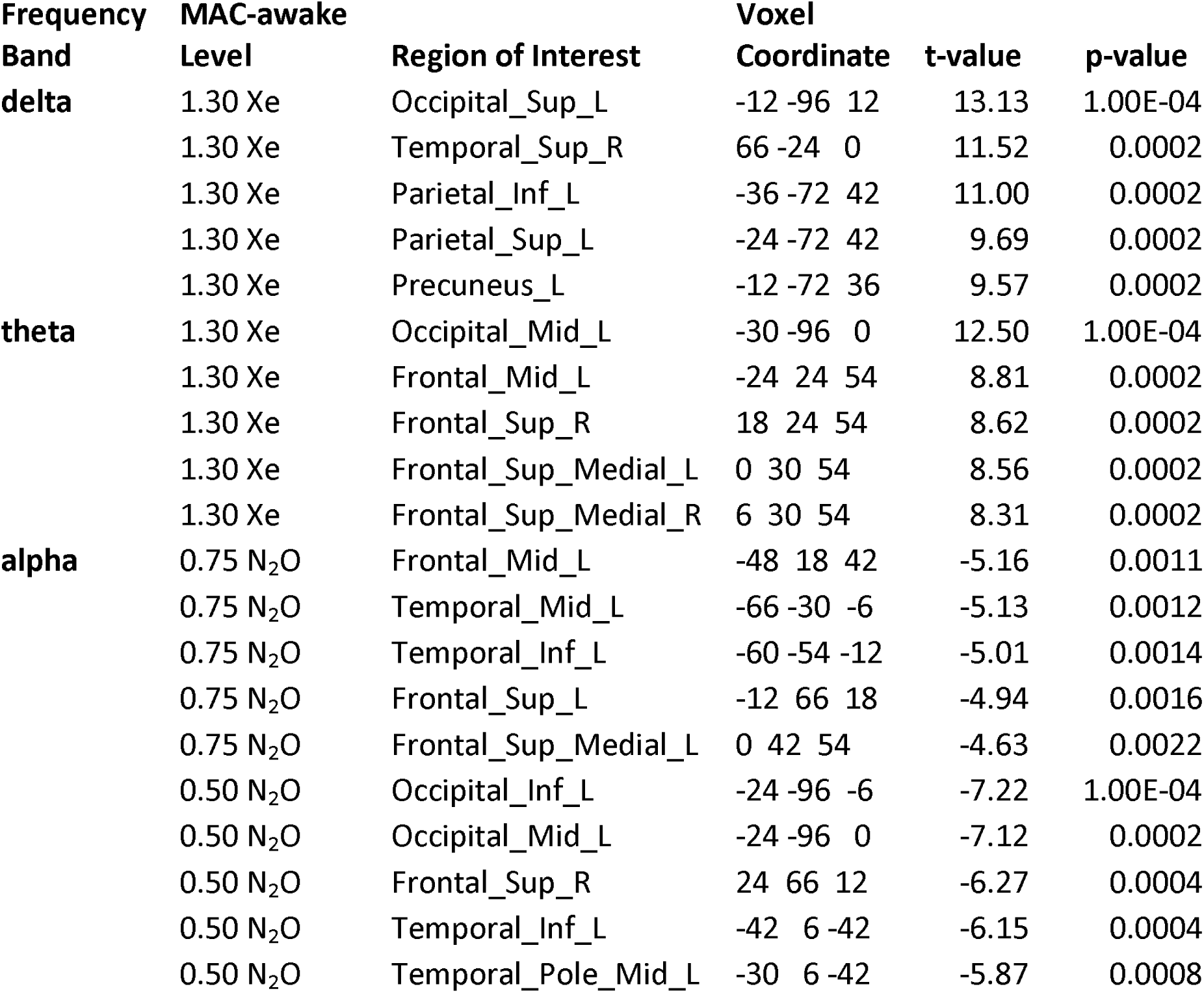
Electroencephalographic sources most significantly altered byXe and N_2_0 administration. Five representative regions of interest were selected from the twenty highest t-value and p-value (p<0.005 for N_2_O; p<0.004 for Xe) regions. Region of interest analysis reveals widespread increases for delta in frontal regions and a widespread theta rise relative to baseline during Xe Loss of Responsiveness (1.30 MAC-awake). The highest N_2_O inspired concentration (0.75 MAC-awake) resulted in primarily frontal alpha reductions while intermediate N_2_O dosage (0.50 MAC-awake) displayed widespread decreases across participants. Voxel coordinates are in AAL atlas coordinate system along with associated labels^46^, [delta (1-4 Hz), theta (4-8 Hz), alpha (8-13 Hz)].

| Frequency Band | MAC-awake Level | Region of Interest | Voxel Coordinate | t-value | p-value |
| --- | --- | --- | --- | --- | --- |
| **delta** | 1.30 Xe | Occipital_Sup_L | -12 -96 12 | 13.13 | 1.00E-04 |
|  | 1.30 Xe | Temporal_Sup_R | 66 -24 0 | 11.52 | 0.0002 |
|  | 1.30 Xe | Parietal_Inf_L | -36 -72 42 | 11.00 | 0.0002 |
|  | 1.30 Xe | Parietal_Sup_L | -24 -72 42 | 9.69 | 0.0002 |
|  | 1.30 Xe | Precuneus_L | -12 -72 36 | 9.57 | 0.0002 |
| **theta** | 1.30 Xe | Occipital_Mid_L | -30 -96 0 | 12.50 | 1.00E-04 |
|  | 1.30 Xe | Frontal_Mid_L | -24 24 54 | 8.81 | 0.0002 |
|  | 1.30 Xe | Frontal_Sup_R | 18 24 54 | 8.62 | 0.0002 |
|  | 1.30 Xe | Frontal_Sup_Medial_L | 0 30 54 | 8.56 | 0.0002 |
|  | 1.30 Xe | Frontal_Sup_Medial_R | 6 30 54 | 8.31 | 0.0002 |
| **alpha** | 0.75 N<sub>2</sub>O | Frontal_Mid_L | -48 18 42 | -5.16 | 0.0011 |
|  | 0.75 N<sub>2</sub>O | Temporal_Mid_L | -66 -30 -6 | -5.13 | 0.0012 |
|  | 0.75 N<sub>2</sub>O | Temporal_Inf_L | -60 -54 -12 | -5.01 | 0.0014 |
|  | 0.75 N<sub>2</sub>O | Frontal_Sup_L | -12 66 18 | -4.94 | 0.0016 |
|  | 0.75 N<sub>2</sub>O | Frontal_Sup_Medial_L | 0 42 54 | -4.63 | 0.0022 |
|  | 0.50 N<sub>2</sub>O | Occipital_Inf_L | -24 -96 -6 | -7.22 | 1.00E-04 |
|  | 0.50 N<sub>2</sub>O | Occipital_Mid_L | -24 -96 0 | -7.12 | 0.0002 |
|  | 0.50 N<sub>2</sub>O | Frontal_Sup_R | 24 66 12 | -6.27 | 0.0004 |
|  | 0.50 N<sub>2</sub>O | Temporal_Inf_L | -42 6 -42 | -6.15 | 0.0004 |
|  | 0.50 N<sub>2</sub>O | Temporal_Pole_Mid_L | -30 6 -42 | -5.87 | 0.0008 |

**Figure 3.**
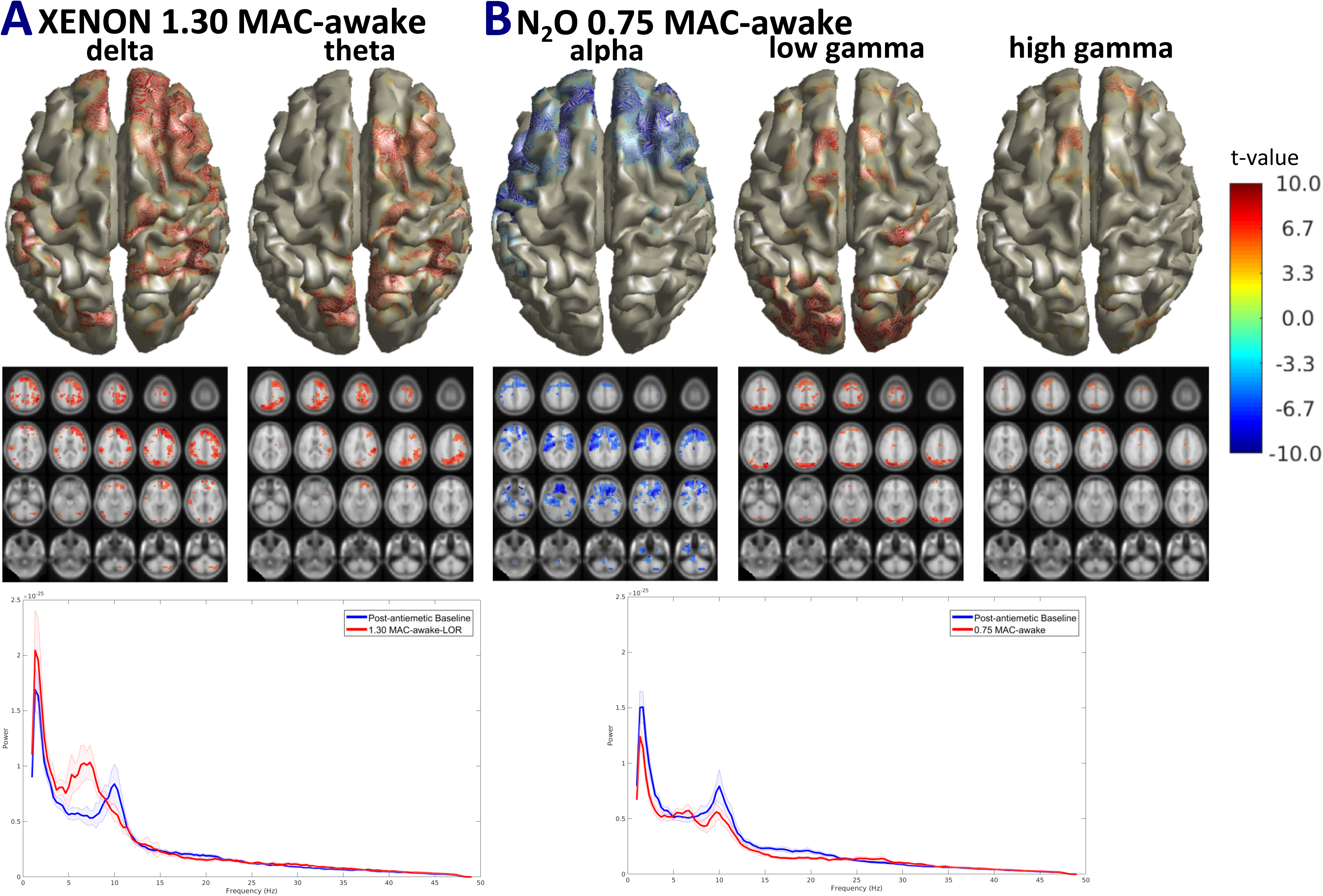
Group level Magnetoencephalography source power for highest administered Xe and N_2_0 concentrations. Maximum statistics and Bonferroni corrected t-maps displaying highly significant (Xe: p=0.004, N_2_O: p=0.005) alterations in magnetoencephalographic sources. Slices display region specific alterations plotted on the template MNI brain. Magnetoencephalography source spectral power (1-49 Hz) is shown for the post-antiemetic baseline and the highest administered doses of the two gases averaged across all magnetoencephalographic sources and all subjects. Shaded regions represent Standard Error. Xe effects (A) at Loss of Responsiveness (1.30 MAC-awake) are increases in low frequency delta (widespread change) and theta activity (right brain lateralized). 0.75 MAC-awake N_2_O (B) results in a primarily frontal alpha reduction and a rise in high frequency gamma activity centered around frontal and occipital regions, [delta (1-4 Hz), theta (4-8 Hz), alpha (8-13 Hz), low gamma (30-49 Hz), high gamma (51-99 Hz)].

For the delta band, subjects that lost responsiveness at Xe 1.3 MAC-awake anesthesia displayed a marked rise in delta band power, primarily involving frontal regions. Theta band power was also significantly increased during Xe Loss of Responsiveness but displayed a different source topology, with widespread changes observed somewhat lateralized to the right hemisphere. Less pronounced effects were observed for 0.75 MAC-awake Xe and were primarily centered on parietal and subcortical regions (see Supplementary Digital Content 2 for details). Conversely, all doses of N_2_O had no substantial effects on low frequency activity. Alpha band power was not affected by Xe but on the other hand, was significantly decreased by N_2_O administration and was predominately frontal in location. Increasing N_2_O concentrations seemed directly correlated to the degree of significance and the spread of alpha reduction (See Supplementary Digital Content 2 for step-wise changes) however, the most pronounced effect was shown at 0.75 MAC-awake N_2_O as demonstrated in figure 3.

Higher frequencies were significantly altered by N_2_O and not by Xe administration, although in this case changes were limited to the highest 0.75 MAC-awake N_2_O concentration given. N_2_O substantially increased both low gamma and high gamma band power and effects were primarily observed in occipital voxels. Finally, under both Xe and N_2_O anesthesia, only a small number of seemingly random voxels seemed to change significantly in the beta band range (see Supplementary Digital Content 2 for details) suggesting that neither gas had a significant effect in this frequency.

#### Electroencephalography Results

Significant power changes in electroencephalographic source power are outlined in Figure 4 with a condensed list of corresponding regions of interest shown in Table 2 (refer to Supplementary Digital Content 2 for complete list of significant regions of interest). Xe inhalation significantly increased low delta and theta band frequency power whereas it did not induce significant changes in higher frequencies. In contrast, N_2_O administration did not alter low frequency delta and theta power. Instead, N_2_O alone reduced frontal alpha power in a dose-dependent manner (see below for details). Neither anesthetic significantly altered high frequency beta or gamma electroencephalographic source power.

**Figure 4.**
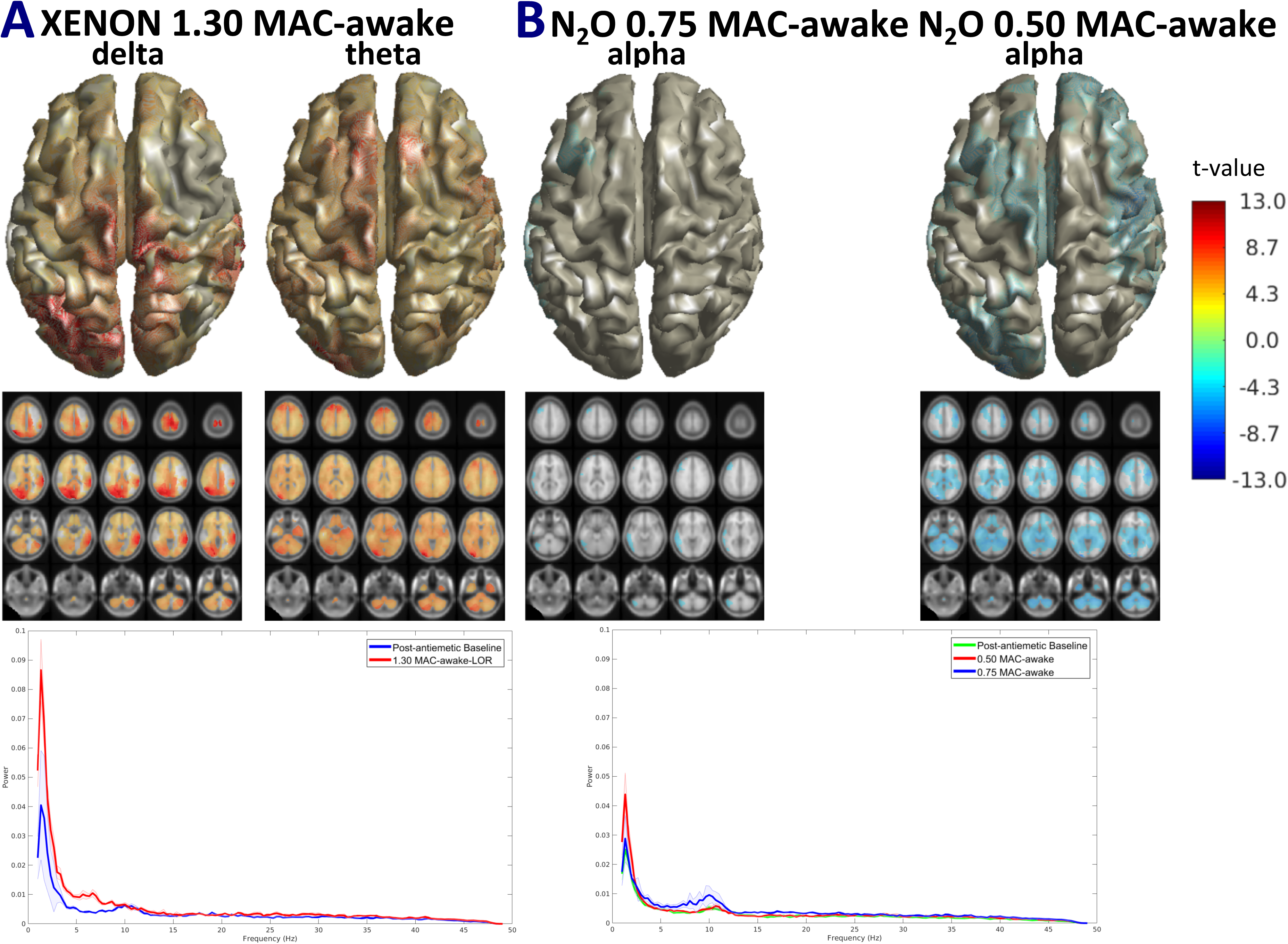
Group level Electroencephalography source power with most significant effects under Xe and N_2_0 anesthesia. Maximum statistics and Bonferroni corrected t-maps displaying highly significant (Xe: p=0.004, N_2_O: p=0.005) alterations in electroencephalographic sources. Slices display region specific alterations plotted on the template MNI brain. Electroencephalography source spectral power (1-49 Hz) is shown for the post-antiemetic baseline and the highest administered doses of the two gases averaged across all electroencephalographic sources and all subjects. Shaded regions represent Standard Error. Loss of Responsiveness at 1.30 MAC-awake Xe administration (A) yields pronounced widespread increases in low frequency delta and theta activity. N_2_O effect (B) is a reduction in alpha activity that is global and more pronounced for 0.50 MAC-awake N_2_O. 0.75 MAC-awake N_2_O effects were subtle and centered on frontal and temporal regions. There were no statistically significant high frequency power changes, [delta (1-4 Hz), theta (4-8 Hz), alpha (8-13 Hz)].

The delta and theta band significant increases observed in the magnetoencephalography power were reflected in the electroencephalography data, though they were widespread and evident only at the highest 1.3 MAC-awake Xe concentration and not low concentrations of Xe or N_2_O. In addition, the localization of alterations was substantially different between the two modalities, particularly in the delta band, where more widespread changes were evident. Like magnetoencephalographic responses, alpha band power was significantly depressed by N_2_O but not Xe and was predominately frontal in topology. Of note, is that for electroencephalographic power, the highest degree of reduction in terms of the number of significant voxels and corresponding t-values, was observed at 0.50 MAC-awake N_2_O while only a small number of frontal and parietal voxels displayed decreased power at 0.75 MAC-awake N_2_O (see Supplementary Digital Content 2 for details).

No significant differences to baseline were observed in higher frequency electroencephalographic power for all beta, low and high gamma ranges.

### Across Gas Statistical Analysis

To look for specific differences between Xe and N_2_O effects, the three equi-MAC-awake concentrations of Xe and N_2_O administered were compared within each equi-MAC-awake level. A figure illustrating group level frequency specific t-statistic maps across the anesthetic agents for both magnetoencephalography and electroencephalography power can be found in Supplementary Digital Content 3.

Unlike the within gas comparison against the post-antiemetic baseline and despite the presence of an effect of gas on the calculated t-maps, most observations did not survive multiple comparisons correction using maximum statistics and Bonferroni correction. The results that demonstrated significant changes are shown in Figure 5 and only relate to alpha band differences between the two gases. First, magnetoencephalographic alpha band source power was significantly lower for 0.75 equi-MAC awake N_2_O against Xe administration in four frontal regions (See Table 3 for details). In addition, electroencephalography source power in the alpha band range was significantly smaller at 50 equi-MAC awake N_2_O against Xe over a number of AAL regions but was primarily frontal (See Table 3 for details and Supplementary Digital Content 3 for full list). Additionally, 4 frontal regions displayed significant reductions in theta power for the 0.50 MAC-awake N_2_O dosage relative to Xe. Minor differences at this equi MAC-awake concentration were observed in delta power and in beta power in a small number of regions (refer to Supplementary Digital Content 3B, Table for further details).

**Table 3.**
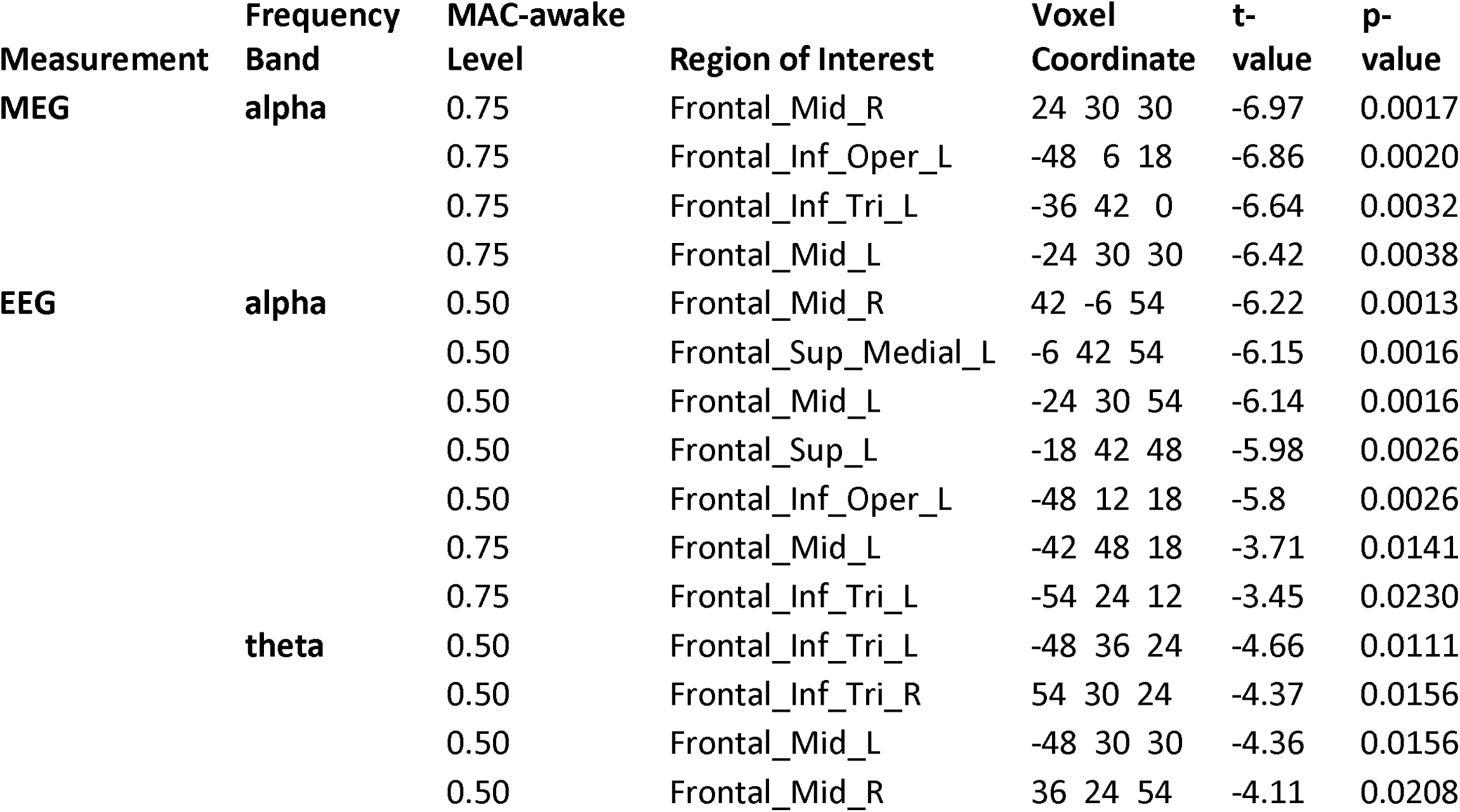
Magnetoencephalographic and Electroencephalographic sources most significantly altered in equivalent gas concentrations of Xe and N_2_0. Representative regions of interest were selected from the twenty highest t-value and p-value (p<0.004) regions. Magnetoencephalographic alpha band power was significantly lower for 0.75 equi-MAC awake N_2_O against Xe administration in four frontal regions). Electroencephalography source power in the alpha band range was significantly smaller at 0.50 equi MAC-awake N_2_O against Xe as well as 0.75 equi MAC-awake and was also primarily frontal. Frontal regions displayed significant reductions in theta power for the 0.50 MAC-awake N_2_O dosage relative to Xe. Voxel coordinates are in AAL atlas coordinate system along with associated labels^46^, [delta (1-4 Hz), theta (4-8 Hz), alpha (8-13 Hz)].

**Figure 5.**
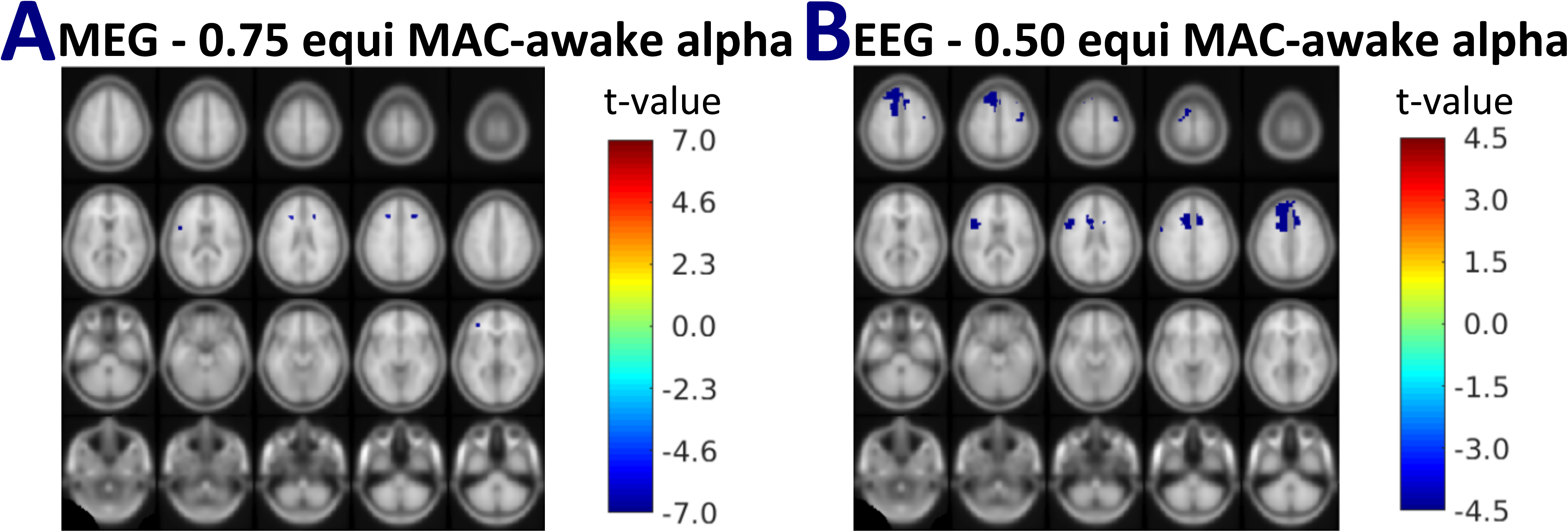
Alpha Magnetoencephalographic and Electroencephalographic source power is substantially less for N_2_0 compared to Xe administration. Maximum statistics and Bonferroni corrected t-maps are plotted on the template MNI brain. Alpha bandwidth magnetoencephalography (MEG - A) source power was considerably lower for N_2_O when compared to Xe anesthesia (0.75 equi MAC-awake) Alpha bandwidth electroencephalographic source power (EEG - B) was substantially less for 0.50 MAC-awake N_2_O against Xe administration and was specific to numerous regions. Representative regions of interest shown here reveal that differences between the two gases were predominately frontal. (AAL Atlas Labels: Frontal_Mid_R, Frontal_Sup_Medial_L, Frontal_Mid_L, Frontal_Sup_L and corresponding Voxel Coordinates). The difference in scale between A and B should be noted, [alpha (8-13 Hz)].

## 4. Discussion

The primary aim of this study was to examine modality and agent dependent dissociative anesthesia variations using electroencephalographic and magnetoencephalographic changes in source power following administration of the gaseous anesthetics Xe and N_2_O in healthy volunteers. Subanesthetic doses of N_2_O yielded no significant effects on low frequency delta and theta activity whereas Xe increased total power in both bands. These findings were reflected in both the magnetoencephalography and electroencephalography data. Alpha power depression was observed only under N_2_O administration and was predominately of frontal origin. Higher frequency activity increases were observed in the magnetoencephalographic but not the electroencephalographic signals for N_2_O, with occipital low gamma and widespread high gamma rise in power. Xe showed no significant gamma band alterations and neither gas gave rise to important beta band activity when compared to baseline. Finally, contrasting the effects of N_2_O relative to Xe did not reflect the differences observed when the agents were evaluated independently against the post-antiemetic baseline. N_2_O showed reduced alpha band magnetoencephalographic source power compared to Xe for the highest inhaled concentration of N_2_O and more pronounced effects for the intermediate dosage administered in electroencephalographic source power. The latter observation is congruent with the within gas electroencephalography results, where alpha changes were more pronounced for this concentration, compared to the highest concentration administered. In summary, electrophysiological findings reveal an array of differences and similarities amongst the two agents that interestingly extend not only to the remnant NMDA-targeting agent ketamine, but also to the GABA-agonists, however, they do not appear to support a universal mechanism for Xe and N_2_O dissociative anesthesia, at least in terms of spectral power.

An important result is the substantial increase in delta power under subanesthetic Xe anesthesia which has been documented in other electrophysiological investigations of Xe ^17,19^. Increases in delta band power are commonly observed in unconsciousness under propofol anesthesia and often demonstrate patterns of increase that are directly correlated to dosage and inversely correlate with participant responsiveness^7-9,50^. Interestingly, the rise in delta may be relevant to Xe’s potentiation effect on depolarization dependent potassium channels^51^ and subsequent hyperpolarization of cortical neurons, which, as has been described for propofol, might result in cortical bistability^9,50^. A recent study comparing Xe, propofol and ketamine subanesthetic dosage using electroencephalographic recordings under Transcranial Magnetic Stimulation attributed the stimulation evoked global negative wave observed under Xe anesthesia to this cortical hyperpolarized state, while propofol-induced local positive-negative waves were thought to be the result of GABA-induced local hyperpolarization^52^. Similarity in the two agents was signified in equivalent, substantially lower than wakefulness values found for a measure of complexity. Conversely, ketamine administration, which like N_2_O has no influence on these molecular targets, resulted in multiple activation waves and a high measure of complexity, comparable to wakefulness, suggestive of no effect on cortico-cortical interactions^52^. In summary, the potentiating effect on potassium currents and resulting generation of delta oscillatory activity with Xe but not N_2_O^51,53^ might offer an explanation as to why we report delta band changes for Xe alone.

Additional lower frequency changes are worth exploring further. The observed theta power increases under Xe anesthesia is in line with the electroencephalography literature^17,19^. The absence of N_2_O-induced theta band changes has been previously reporded^54^, however remains unlike most reports pointing to theta power increases under N_2_O^15,16^ and ketamine^12,32,55,56^. The significant reduction in electroencephalographic alpha power upon subanesthetic N_2_O administration has been observed^13,54^ and replicated using ketamine anesthetic dosage^56^. Importantly, due to the close resemblance to our paradigm, a theta rise was replicated using magnetoencephalography and subanesthetic ketamine doses^32^. Intriguingly, we do not report the same result for electroencephalography and magnetoencephalography power. This may be explained by the necessary common average re-referencing scheme^57^ performed only on the electroencephalography data and used as the standard re-referencing scheme for EEG source imaging^29^. It could alternatively relate to dosage, with previous work reporting no alterations in alpha at lower concentrations (40 % N_2_O/O_2_^15^; 47 % in our study) but significant reductions at higher concentrations (60 % N_2_O/O_2_^54^,not administered in our study due to safety concerns). Xe did not significantly alter alpha power, as has been formerly presented^17^. A general point to be made here is that observed power changes, can be complemented by functional connectivity measures which may yield more powerful functional specificity^56^ (such investigations will be the subject of future work). The distinctive effects of the two agents on the theta and alpha band we outlined by contrasting their anesthetic action on each individual, may ultimately relate to large-scale (low frequency dependent) network alterations along the anterior to posterior axis, but further investigations are required to consider this postulation.

Higher frequency increases in the gamma range as described here are common under N_2_O-induced reductions in consciousness^13,14^. The widespread high gamma power rise reported additionally resembles global gamma power increases in magnetoencephalographic recordings under ketamine infusion^32,58^. Furthermore, we report no significant changes in beta power, as has been previously shown for lower N_2_O administered doses^15,16^ and for GABA-ergic anesthetics^50,59^. Xe inhalation did not exhibit significant effects on high frequency gamma and beta activity and previous findings remain unclear on this relationship^17,20^. Nonetheless, the different high frequency profiles observed for the two agents are of particular interest, as they may be suggestive of N_2_O mediated noise, influencing the firing of pyramidal neurons and hence disrupting consciousness^60^.

On a different note, important points emerge pertaining to the protocol followed. Noteworthy is the fact that high frequency N_2_O observations were not replicated for Xe, which is of particular interest in magnetoencephalography datasets. Electromagnetic gamma activity is often contaminated with muscle artefacts which commonly oscillate in a similar range of frequencies^61^, a critical concern in subanesthetic gaseous anesthesia, where psychomotor agitation is pronounced, particularly reflected in increased jaw movements^30^. Since changes in gamma were not evident in both gas conditions for the same volunteers, and considering that rigorous measures were taken to eliminate head and body movement, we can assume true anesthetic induced gamma band variations. In addition, high frequency results demonstrate significant power changes for magnetoencephalography but not electroencephalography data. This may be credited to the wider frequency range of acquisition that the considered magnetoencephalography data possesses, which extends to gamma activity^62^. On a similar point, overall differences observed across the two modalities may be attributed to the higher spatial and source localization accuracy of magnetoencephalographic recordings^29,63,64^. We therefore speculate that magnetoencephalography (and simultaneous magnetoencephalography/electroencephalography) source imaging may transpire as an excellent alternative to the more commonly employed electroencephalography, for the investigation of the fine-tuned and region specific neural networks involved in reductions of consciousness under anesthesia and beyond.

Various additional strengths of this study are worth mentioning. The cross-over design applied here enabled the true and direct comparison of the two administered agents and induced reductions in consciousness. In addition, the fact that various equi-MAC-awake concentrations of Xe and N_2_O were administered with the same gas wash-in, maintenance and wash-out steps being taken at each, reduced any effects that lower concentrations may have and therefore increased the purity of the comparison. In addition, the administration outcomes could only be credited to each gaseous anesthetic as no other psychoactive medication was administered; a common confounding factor in anesthesia research. The use of magnetoencephalography and high density electroencephalography yielded high-resolution recordings hence, allowing for source level imaging techniques to be applied, in order to reveal the region specific origins of any observed signals.

On the contrary, there are multiple important weaknesses to mention. Firstly, the direct comparison of the magnetoencephalography and electroencephalography results is rendered difficult, due to a number of differences in the datasets and analysis pipeline. Participant numbers and sampling rates were significantly less for electroencephalographic data, and sample sizes have shown a contribution to the quality of source localization^65^. Necessary re-referencing was performed on electroencephalograms, while magnetoencephalographic recordings alone were filtered using temporal signal source space separation. Also, forward model calculation significantly differs for the two modalities since electrical but not electromagnetic signals are influenced by the skull^29^. With this volume conduction issue in mind, sensor level data have not been demonstrated here. Despite the above, combining the two datasets prior to beamforming was not favored due to associated issues described in the literature^66^ and our interest in modality dependent changes. No connectivity analyses were performed to investigate network changes under anesthesia. A final drawback, was the use of highly conservative statistics that resulted in the loss of potentially interesting information.

In conclusion, this study investigated the effects of increasing equivalent doses of Xe and N_2_O on magnetoencephalographic and electroencephalographic source power in healthy volunteers. Magnetoencephalography and electroencephalography recordings revealed increased low frequency delta and theta power at the highest Xe concentration and reduced alpha power under subanesthetic N_2_O administration. Finally, a rise in high frequency gamma activity was demonstrated in the magnetoencephalography data. Higher specificity and range of magnetoencephalography results, point to important benefits of exploiting this underused technique to enhance temporally and spatially specific imaging. Power results provide novel information on the mechanisms underlying reductions in consciousness resulting from these unconventional NMDA-antagonist gaseous anesthetics.

## Supporting information

Supplementary Digital Content 1

Supplementary Digital Content 2

Supplementary Digital Content 3

## Acknowledgments

The authors acknowledge the facilities, and the scientific and technical assistance of the National Imaging Facility at the Magnetic Resonance Imaging and Magnetoencephalography Units, Swinburne Neuroimaging Facility, Swinburne University of Technology. We thank Richard Aveyard Ph.D., Imaging IT Coordinator, Swinburne Neuroimaging Facility, Swinburne University of Technology for assistance with computational issues and for valuable ideas. Many thanks to Heonsoo Lee Ph.D. and Uncheol Lee Ph.D. from the Department of Anesthesiology, University of Michigan Medical School, Ann Arbor, Ml, United States; Center for Consciousness Science, University of Michigan Medical School, Ann Arbor, Ml, United States for many stimulating discussions on analysis, consciousness and beyond throughout the study.

## Abbreviated Title

Xe: Source power changes
N_2_O: anesthesia

