## Supplementary Digital Content 1 for "In search of Universal Cortical Power Changes Linked to NMDA-Antagonist based Anesthetic Induced Reductions in Consciousness"

The datasets utilized in the experiment as well as preliminary power results are summarized below to demonstrate differences between the datasets used and resulting power analysis.

*Supplementary Table 1A. Number of participant datasets used in statistical analysis.* Table demonstrates number of source reconstructed images utilized in statistical analysis and the number of samples used in each subject to generate them.

| Dataset Type | Resting Baseline | Pre-antiemetic Baseline | Post-antiemetic Baseline | 0.25 MAC-awake | 0.50  MAC-awake | 0.75 MAC-awake | 1.3  MAC-awake  (LOR) |
| --- | --- | --- | --- | --- | --- | --- | --- |
| MEG Xe (51000 samples) | 21 | 21 | 21 | 21 | 21 | 20 | 16 |
| MEG N_2_O  (51000 samples) | 21 | 21 | 21 | 21 | 21 | 21 | N/A |
| EEG Xe  (13824 samples) | 18 | 19 | 19 | 19 | 18 | 18 | 14 |
| EEG N_2_O  (13824 samples) | 18 | 18 | 19 | 18 | 18 | 18 | N/A |

*Supp. Table 1B. Total power at each baseline and inspired gas concentration and change relative to post-antiemetic baseline.* Power changes with ±1 SD for magnetoencephalography (MEG) and electroencephalography (EEG) data in each band for the post- antiemetic baseline and increasing Xe and N_2_O concentrations are shown in the first and second section of the table and are in T^2^ for MEG and in μV^2^ for EEG. Relative change to post-antiemetic baseline ([(Gas power-Baseline power)/Baseline power] x 100 in %) are shown in the third and fourth section of the table.

| **TOTAL POWER**  **CHANGES** | |  | |  |  |  |  |  |  |
| --- | --- | --- | --- | --- | --- | --- | --- | --- | --- |
| **MEG** |  |  | |  |  |  |  |  |  |
| **Frequency Band** | **Gas** | | **Resting Baseline** | **Pre-antiemetic Baseline** | **Post-antiemetic Baseline** | **0.25 MAC-awake** | **0.50 MAC-awake** | **0.75 MAC-awake** | **1.30 MAC-awake (LOR)** |
| **delta** | **Xe** | 2.04E-25 | | 2.21E-25 | 2.71E-25 | 2.85E-25 | 2.94E-25 | 3.58E-25 | 7.10E-25 |
|  | ±1 SD | 2.46E-26 | | 2.93E-26 | 2.70E-26 | 4.80E-26 | 4.02E-26 | 4.25E-26 | 3.36E-25 |
|  | **N2O** | 1.86E-25 | | 2.13E-25 | 2.55E-25 | 2.34E-25 | 2.48E-25 | 2.80E-25 |  |
|  | ±1 SD | 1.96E-26 | | 2.71E-26 | 3.68E-26 | 3.15E-26 | 2.82E-26 | 5.33E-26 |  |
| **theta** | **Xe** | 1.57E-25 | | 1.64E-25 | 1.89E-25 | 1.88E-25 | 1.89E-25 | 2.23E-25 | 3.49E-25 |
|  | ±1 SD | 1.75E-26 | | 1.96E-26 | 1.94E-26 | 1.83E-26 | 1.83E-26 | 2.14E-26 | 9.47E-26 |
|  | **N2O** | 1.42E-25 | | 1.45E-25 | 1.70E-25 | 1.56E-25 | 1.58E-25 | 1.77E-25 |  |
|  | ±1 SD | 1.38E-26 | | 1.46E-26 | 1.78E-26 | 1.54E-26 | 1.49E-26 | 4.85E-26 |  |
| **alpha** | **Xe** | 3.64E-25 | | 3.80E-25 | 4.33E-25 | 3.71E-25 | 3.30E-25 | 3.57E-25 | 2.82E-25 |
|  | ±1 SD | 3.68E-26 | | 4.47E-26 | 3.98E-26 | 3.49E-26 | 3.01E-26 | 3.77E-26 | 3.16E-26 |
|  | **N2O** | 3.60E-25 | | 3.67E-25 | 4.28E-25 | 3.56E-25 | 3.01E-25 | 2.39E-25 |  |
|  | ±1 SD | 3.38E-26 | | 3.78E-26 | 3.94E-26 | 3.61E-26 | 3.16E-26 | 2.98E-26 |  |
| **beta** | **Xe** | 2.26E-25 | | 2.36E-25 | 2.71E-25 | 2.75E-25 | 2.72E-25 | 3.20E-25 | 3.44E-25 |
|  | ±1 SD | 1.52E-26 | | 1.60E-26 | 1.41E-26 | 1.94E-26 | 1.49E-26 | 2.22E-26 | 2.39E-26 |
|  | **N2O** | 2.17E-25 | | 2.26E-25 | 2.59E-25 | 2.62E-25 | 2.64E-25 | 2.62E-25 |  |
|  | ±1 SD | 1.22E-26 | | 1.96E-26 | 1.40E-26 | 1.52E-26 | 1.72E-26 | 1.67E-26 |  |
| **low gamma** | **Xe** | 1.05E-25 | | 1.15E-25 | 1.51E-25 | 1.61E-25 | 1.65E-25 | 1.83E-25 | 2.16E-25 |
|  | ±1 SD | 4.54E-27 | | 5.19E-27 | 1.33E-26 | 2.69E-26 | 1.58E-26 | 1.01E-26 | 1.57E-26 |
|  | **N2O** | 1.08E-25 | | 1.23E-25 | 1.35E-25 | 1.51E-25 | 1.55E-25 | 1.75E-25 |  |
|  | ±1 SD | 4.56E-27 | | 3.17E-26 | 6.49E-27 | 1.03E-26 | 8.31E-27 | 1.08E-26 |  |
| **high gamma** | **Xe** | 1.98E-25 | | 2.26E-25 | 3.10E-25 | 3.29E-25 | 3.23E-25 | 3.45E-25 | 4.13E-25 |
|  | ±1 SD | 6.91E-27 | | 9.36E-27 | 4.42E-26 | 1.27E-25 | 6.79E-26 | 2.59E-26 | 3.36E-26 |
|  | **N2O** | 2.15E-25 | | 3.33E-25 | 2.67E-25 | 2.86E-25 | 3.02E-25 | 3.50E-25 |  |
|  | ±1 SD | 6.01E-27 | | 3.33E-26 | 1.18E-26 | 2.15E-26 | 1.59E-26 | 4.84E-26 |  |
| **EEG** |  |  | |  |  |  |  |  |  |
| **Frequency Band** | **Gas** | **Resting Baseline** | | **Pre-antiemetic Baseline** | **Post-antiemetic Baseline** | **0.25 MAC-awake** | **0.50 MAC-awake** | **0.75 MAC-awake** | **1.30 MAC-awake (LOR)** |
| **delta** | **Xe** | 0.05 | | 0.05 | 0.09 | 0.11 | 0.18 | 0.20 | 0.43 |
|  | ±1 SD | 0.01 | | 0.00 | 0.04 | 0.04 | 0.07 | 0.08 | 0.13 |
|  | **N2O** | 0.05 | | 0.07 | 0.08 | 0.12 | 0.13 | 0.29 |  |
|  | ±1 SD | 0.00 | | 0.02 | 0.02 | 0.04 | 0.04 | 0.11 |  |
| **theta** | **Xe** | 0.03 | | 0.03 | 0.08 | 0.05 | 0.09 | 0.08 | 0.22 |
|  | ±1 SD | 0.00 | | 0.00 | 0.09 | 0.01 | 0.06 | 0.06 | 0.15 |
|  | **N2O** | 0.03 | | 0.04 | 0.06 | 0.05 | 0.04 | 0.08 |  |
|  | ±1 SD | 0.00 | | 0.00 | 0.04 | 0.02 | 0.01 | 0.04 |  |
| **alpha** | **Xe** | 0.08 | | 0.07 | 0.13 | 0.08 | 0.11 | 0.10 | 0.26 |
|  | ±1 SD | 0.01 | | 0.01 | 0.05 | 0.01 | 0.05 | 0.05 | 0.25 |
|  | **N2O** | 0.08 | | 0.10 | 0.12 | 0.10 | 0.06 | 0.11 |  |
|  | ±1 SD | 0.01 | | 0.02 | 0.01 | 0.02 | 0.01 | 0.08 |  |
| **beta** | **Xe** | 0.09 | | 0.08 | 0.10 | 0.10 | 0.14 | 0.13 | 0.21 |
|  | ±1 SD | 0.01 | | 0.01 | 0.01 | 0.01 | 0.07 | 0.04 | 0.11 |
|  | **N2O** | 0.09 | | 0.10 | 0.11 | 0.13 | 0.10 | 0.15 |  |
|  | ±1 SD | 0.00 | | 0.01 | 0.02 | 0.05 | 0.02 | 0.08 |  |
| **low gamma** | **Xe** | 0.06 | | 0.06 | 0.08 | 0.07 | 0.11 | 0.10 | 0.18 |
|  | ±1 SD | 0.01 | | 0.00 | 0.01 | 0.01 | 0.05 | 0.05 | 0.11 |
|  | **N2O** | 0.06 | | 0.07 | 0.08 | 0.09 | 0.07 | 0.11 |  |
|  | ±1 SD | 0.00 | | 0.01 | 0.01 | 0.04 | 0.01 | 0.06 |  |
| **high gamma** | **Xe** | 0.13 | | 0.13 | 0.18 | 0.14 | 0.17 | 0.16 | 0.26 |
|  | ±1 SD | 0.01 | | 0.02 | 0.04 | 0.02 | 0.05 | 0.03 | 0.12 |
|  | **N2O** | 0.13 | | 0.15 | 0.16 | 0.16 | 0.15 | 0.21 |  |
|  | ±1 SD | 0.00 | | 0.03 | 0.02 | 0.07 | 0.05 | 0.08 |  |
| **RELATIVE POWER CHANGES** | | | |  |  |  |  |  |  |
| **MEG** |  |  | |  |  |  |  |  |  |
| **Frequency Band** | **Gas** | **Resting Baseline** | | **Pre-antiemetic Baseline** | **Post-antiemetic Baseline** | **0.25 MAC-awake** | **0.50 MAC-awake** | **0.75 MAC-awake** | **1.30 MAC-awake (LOR)** |
| **delta** | **Xe** | -20.93 | | -15.51 |  | 10.37 | 22.22 | 61.60 | 208.32 |
|  | ±1 SD | 11.71 | | 11.64 |  | 9.39 | 19.69 | 45.94 | 136.34 |
|  | **N2O** | -36.60 | | -25.59 |  | -4.40 | 7.59 | 19.86 |  |
|  | ±1 SD | 14.03 | | 14.37 |  | 6.37 | 13.37 | 17.84 |  |
| **theta** | **Xe** | -11.94 | | -9.28 |  | 5.81 | 13.24 | 45.32 | 107.43 |
|  | ±1 SD | 11.00 | | 10.86 |  | 8.09 | 16.31 | 41.65 | 82.45 |
|  | **N2O** | -26.97 | | -22.47 |  | -3.17 | 0.38 | 12.38 |  |
|  | ±1 SD | 14.21 | | 12.76 |  | 6.23 | 11.84 | 16.96 |  |
| **alpha** | **Xe** | -19.68 | | -15.24 |  | -13.77 | -22.42 | -16.79 | -44.59 |
|  | ±1 SD | 11.14 | | 9.79 |  | 6.85 | 10.01 | 9.36 | 12.93 |
|  | **N2O** | -18.75 | | -17.15 |  | -15.76 | -28.88 | -44.36 |  |
|  | ±1 SD | 9.28 | | 8.83 |  | 6.98 | 11.18 | 12.64 |  |
| **beta** | **Xe** | -14.25 | | -8.79 |  | 7.62 | 13.58 | 44.00 | 49.04 |
|  | ±1 SD | 11.14 | | 10.41 |  | 9.05 | 16.04 | 37.62 | 54.15 |
|  | **N2O** | -27.24 | | -23.15 |  | 7.15 | 16.14 | 15.64 |  |
|  | ±1 SD | 15.58 | | 15.31 |  | 8.77 | 20.41 | 20.77 |  |
| **low gamma** | **Xe** | -26.16 | | -16.76 |  | 11.29 | 23.83 | 53.32 | 77.50 |
|  | ±1 SD | 13.34 | | 14.82 |  | 8.66 | 19.18 | 47.03 | 89.86 |
|  | **N2O** | -32.19 | | -22.40 |  | 18.14 | 28.58 | 46.27 |  |
|  | ±1 SD | 17.59 | | 18.05 |  | 11.11 | 20.39 | 25.59 |  |
| **high gamma** | **Xe** | -33.83 | | -23.41 |  | 10.31 | 15.04 | 37.23 | 54.98 |
|  | ±1 SD | 13.73 | | 13.36 |  | 9.17 | 13.74 | 38.50 | 71.69 |
|  | **N2O** | -29.81 | | -1.48 |  | 14.02 | 25.09 | 45.66 |  |
|  | ±1 SD | 16.07 | | 29.89 |  | 9.87 | 18.48 | 24.49 |  |
| **EEG** |  |  | |  |  |  |  |  |  |
| **Frequency Band** | **Gas** | **Resting Baseline** | | **Pre-antiemetic Baseline** | **Post-antiemetic Baseline** | **0.25 MAC-awake** | **0.50 MAC-awake** | **0.75 MAC-awake** | **1.30 MAC-awake (LOR)** |
| **delta** | **Xe** | -46.77 | | -39.52 |  | 19.24 | 88.38 | 118.59 | 260.30 |
|  | ±1 SD | 5.56 | | 5.81 |  | 18.25 | 36.89 | 38.75 | 60.00 |
|  | **N2O** | -35.03 | | -16.01 |  | 48.85 | 73.27 | 269.65 |  |
|  | ±1 SD | 9.18 | | 11.38 |  | 30.83 | 36.93 | 92.54 |  |
| **theta** | **Xe** | -58.85 | | -54.65 |  | -34.58 | 9.34 | 2.10 | 116.28 |
|  | ±1 SD | 9.00 | | 10.16 |  | 14.50 | 20.51 | 20.13 | 75.75 |
|  | **N2O** | -40.44 | | -30.33 |  | -10.99 | -22.09 | 40.96 |  |
|  | ±1 SD | 7.85 | | 9.41 |  | 11.21 | 10.80 | 30.95 |  |
| **alpha** | **Xe** | -40.59 | | -42.74 |  | -36.21 | -20.52 | -30.36 | 49.25 |
|  | ±1 SD | 7.11 | | 5.96 |  | 7.01 | 8.58 | 14.43 | 30.88 |
|  | **N2O** | -32.28 | | -10.03 |  | -12.14 | -44.02 | -5.71 |  |
|  | ±1 SD | 8.97 | | 6.47 |  | 6.92 | 8.10 | 9.05 |  |
| **beta** | **Xe** | -17.63 | | -17.81 |  | -2.02 | 30.82 | 17.18 | 49.65 |
|  | ±1 SD | 3.01 | | 3.69 |  | 3.14 | 11.21 | 13.41 | 26.10 |
|  | **N2O** | -17.30 | | -9.44 |  | 13.11 | -12.94 | 34.11 |  |
|  | ±1 SD | 5.07 | | 5.48 |  | 11.75 | 6.77 | 22.00 |  |
| **low gamma** | **Xe** | -22.01 | | -22.52 |  | -9.88 | 28.37 | 18.47 | 73.95 |
|  | ±1 SD | 4.06 | | 5.06 |  | 4.30 | 10.10 | 33.71 | 32.58 |
|  | **N2O** | -22.46 | | -15.05 |  | 13.68 | -14.18 | 32.32 |  |
|  | ±1 SD | 4.93 | | 5.43 |  | 17.90 | 5.38 | 20.27 |  |
| **high gamma** | **Xe** | -30.36 | | -28.06 |  | -20.50 | -7.93 | -16.26 | 9.09 |
|  | ±1 SD | 6.18 | | 10.45 |  | 6.39 | 6.51 | 7.88 | 11.57 |
|  | **N2O** | -20.96 | | -5.85 |  | -1.21 | -6.06 | 28.56 |  |
|  | ±1 SD | 6.18 | | 5.22 |  | 6.09 | 5.38 | 19.69 |  |
