## Supplementary Digital Content 2 for "In search of Universal Cortical Power Changes Linked to NMDA-Antagonist based Anesthetic Induced Reductions in Consciousness"

Significant maximum statistics (p=0.025) corrected t-statistic maps of the power changes relative to baseline across subjects that demonstrate trends in the data with increasing gas concentration [0.25, 0.50, 0.75 MAC-awake for Xe, N_2_O and 1.30 MAC-awake (Loss of Responsiveness) for Xe] in magnetoencephalography and electroencephalography datasets are shown in Supp. Figure 2.

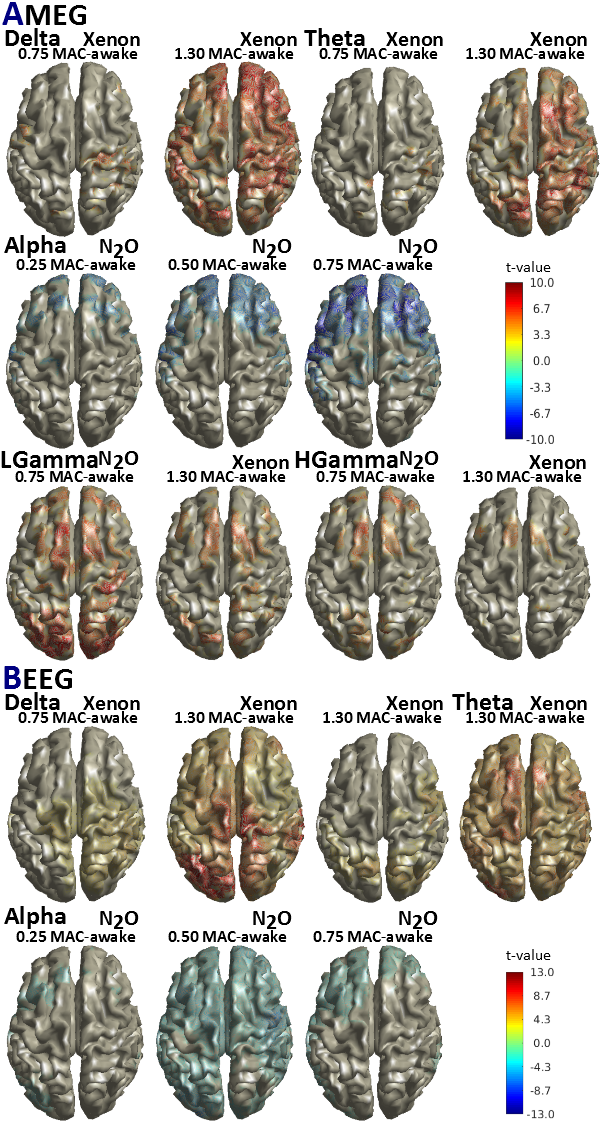

*Supp. Figure 2.* *Group level source power t-statistic maps under increasing doses of Xe and N_2_O relative to post-antiemetic baseline (baseline 3).* Maximum statistics corrected (without subsequent Bonferroni correction) t-values reveal significant (p<0.025) changes in the delta, theta and alpha bands in both magnetoencephalogram (A) and electroencephalogram (B) results for different gas concentrations (0.25, 0.50, 0.75, 1.30 MAC-awake). High frequency power (Lgamma, Hgamma) is only significantly altered for MEG data and only for the highest administered gas concentrations. The difference in scale between A and B should be noted. [delta (1-4 Hz), theta (4-8 Hz), alpha (8-13 Hz), Lgamma: low gamma (30-49 Hz), Hgamma: high gamma (51-99 Hz)].

Highly significant (Xe: p=0.004 ; N_2_O: p=0.005) power changes for the two gases in increasing gas levels contrasted to the post-antiemetic baseline reveal region specific changes in each frequency band investigate. Supp. Tables 2A and 2B give a full account of all significantly altered regions relative to baseline.

*Supp. Table 2A. Magnetoencephalographic sources most significantly altered by Xe and N_2_O administration.* Significantly (p<0.005 for N_2_O; p<0.004 for Xe) altered regions relative to post-antiemetic baseline for each frequency band. Voxel coordinates are in AAL atlas coordinate system along with associated labels^43^. [delta (1-4 Hz), theta (4-8 Hz), alpha (8-13 Hz), low gamma (30-49 Hz), high gamma (51-99 Hz)].

|  | **Delta**  **MAC-awake Level** | | **Region of Interest** | | | | **Voxel Coordinate** | | | | | | **t-value** | | **p-value** | | | | |
| --- | --- | --- | --- | --- | --- | --- | --- | --- | --- | --- | --- | --- | --- | --- | --- | --- | --- | --- | --- |
|  | 1.30 Xenon | | Parietal_Inf_R | | | | 54 -42 48 | | | | | | 9.61 | | 0.0001 | | | | |
|  |  | | Precentral_R | | | | 54 6 42 | | | | | | 9.36 | | 0.0002 | | | | |
|  |  | | Cingulum_Ant_R | | | | 6 48 24 | | | | | | 9.36 | | 0.0002 | | | | |
|  |  | | Frontal_Sup_Medial_L | | | | 0 54 24 | | | | | | 9.25 | | 0.0002 | | | | |
|  |  | | Frontal_Sup_Medial_R | | | | 12 48 24 | | | | | | 9.13 | | 0.0002 | | | | |
|  |  | | Precuneus_L | | | | 0 -48 54 | | | | | | 9.13 | | 0.0002 | | | | |
|  |  | | Frontal_Sup_R | | | | 18 54 30 | | | | | | 9.11 | | 0.0002 | | | | |
|  |  | | Fusiform_R | | | | 48 -66 -18 | | | | | | 8.82 | | 0.0002 | | | | |
|  |  | | Frontal_Mid_R | | | | 48 24 36 | | | | | | 8.80 | | 0.0002 | | | | |
|  |  | | Temporal_Inf_R | | | | 48 -66 -12 | | | | | | 8.47 | | 0.0002 | | | | |
|  |  | | Temporal_Inf_L | | | | -54 -60 -12 | | | | | | 8.45 | | 0.0002 | | | | |
|  |  | | Cuneus_L | | | | -18 -72 36 | | | | | | 8.34 | | 0.0002 | | | | |
|  |  | | SupraMarginal_L | | | | -54 -30 24 | | | | | | 8.32 | | 0.0002 | | | | |
|  |  | | Cingulum_Mid_R | | | | 6 42 30 | | | | | | 8.27 | | 0.0002 | | | | |
|  |  | | Frontal_Inf_Oper_R | | | | 54 12 12 | | | | | | 8.13 | | 0.0002 | | | | |
|  |  | | Frontal_Sup_L | | | | -12 48 42 | | | | | | 8.08 | | 0.0002 | | | | |
|  |  | | Postcentral_L | | | | -60 -18 30 | | | | | | 8.08 | | 0.0002 | | | | |
|  |  | | Frontal_Med_Orb_R | | | | 6 54 -6 | | | | | | 7.92 | | 0.0004 | | | | |
|  |  | | Precuneus_R | | | | 6 -48 54 | | | | | | 7.88 | | 0.0004 | | | | |
|  |  | | SupraMarginal_R | | | | 42 -42 42 | | | | | | 7.85 | | 0.0004 | | | | |
|  |  | | Occipital_Mid_L | | | | -24 -72 30 | | | | | | 7.76 | | 0.0004 | | | | |
|  |  | | Postcentral_R | | | | 42 -30 60 | | | | | | 7.73 | | 0.0004 | | | | |
|  |  | | Precentral_L | | | | -48 -6 42 | | | | | | 7.71 | | 0.0004 | | | | |
|  |  | | Parietal_Inf_L | | | | -54 -36 42 | | | | | | 7.65 | | 0.0004 | | | | |
|  |  | | Parietal_Sup_R | | | | 30 -48 60 | | | | | | 7.61 | | 0.0004 | | | | |
|  |  | | Frontal_Inf_Tri_R | | | | 48 24 30 | | | | | | 7.60 | | 0.0004 | | | | |
|  |  | | Paracentral_Lobule_R | | | | 12 -30 66 | | | | | | 7.60 | | 0.0004 | | | | |
|  |  | | Occipital_Sup_L | | | | -24 -72 36 | | | | | | 7.49 | | 0.0004 | | | | |
|  |  | | Occipital_Inf_L | | | | -24 -96 -6 | | | | | | 7.43 | | 0.0004 | | | | |
|  |  | | Frontal_Mid_Orb_R | | | | 36 48 -6 | | | | | | 7.41 | | 0.0004 | | | | |
|  |  | | Thalamus_R | | | | 6 -18 6 | | | | | | 7.31 | | 0.0004 | | | | |
|  |  | | Supp_Motor_Area_R | | | | 12 -18 66 | | | | | | 7.28 | | 0.0004 | | | | |
|  |  | | Angular_R | | | | 54 -54 36 | | | | | | 7.18 | | 0.0004 | | | | |
|  |  | | Cingulum_Ant_L | | | | 0 36 0 | | | | | | 7.17 | | 0.0004 | | | | |
|  |  | | Frontal_Med_Orb_L | | | | 0 60 -6 | | | | | | 7.13 | | 0.0004 | | | | |
|  |  | | Temporal_Sup_L | | | | -54 -30 18 | | | | | | 7.03 | | 0.0006 | | | | |
|  |  | | Frontal_Inf_Oper_L | | | | -54 6 12 | | | | | | 7.02 | | 0.0006 | | | | |
|  |  | | Occipital_Inf_R | | | | 42 -66 -12 | | | | | | 7.00 | | 0.0006 | | | | |
|  |  | | Occipital_Mid_R | | | | 30 -84 36 | | | | | | 6.98 | | 0.0006 | | | | |
|  |  | | Calcarine_L | | | | -18 -102 -6 | | | | | | 6.94 | | 0.0006 | | | | |
|  |  | | Supp_Motor_Area_L | | | | -12 6 66 | | | | | | 6.90 | | 0.0006 | | | | |
|  |  | | Rolandic_Oper_R | | | | 60 -18 18 | | | | | | 6.89 | | 0.0006 | | | | |
|  |  | | Temporal_Mid_L | | | | -48 -66 6 | | | | | | 6.84 | | 0.0006 | | | | |
|  |  | | Frontal_Sup_Orb_L | | | | -18 42 -18 | | | | | | 6.82 | | 0.0006 | | | | |
|  |  | | Calcarine_R | | | | 18 -90 0 | | | | | | 6.79 | | 0.0008 | | | | |
|  |  | | Occipital_Sup_R | | | | 24 -84 30 | | | | | | 6.75 | | 0.0010 | | | | |
|  |  | | Paracentral_Lobule_L | | | | -6 -30 54 | | | | | | 6.74 | | 0.0010 | | | | |
|  |  | | Parietal_Sup_L | | | | -24 -72 42 | | | | | | 6.72 | | 0.0010 | | | | |
|  |  | | Temporal_Sup_R | | | | 54 -30 18 | | | | | | 6.64 | | 0.0014 | | | | |
|  |  | | Fusiform_L | | | | -42 -72 -18 | | | | | | 6.61 | | 0.0014 | | | | |
|  |  | | Frontal_Inf_Orb_R | | | | 30 36 -12 | | | | | | 6.57 | | 0.0014 | | | | |
|  |  | | Thalamus_L | | | | -6 -12 12 | | | | | | 6.57 | | 0.0014 | | | | |
|  |  | | Cingulum_Mid_L | | | | 0 -42 48 | | | | | | 6.56 | | 0.0014 | | | | |
|  |  | | Frontal_Mid_Orb_L | | | | -42 54 -6 | | | | | | 6.53 | | 0.0016 | | | | |
|  |  | | Rolandic_Oper_L | | | | -48 -24 18 | | | | | | 6.45 | | 0.0016 | | | | |
|  |  | | Frontal_Inf_Orb_L | | | | -42 42 -12 | | | | | | 6.43 | | 0.0016 | | | | |
|  |  | | Cuneus_R | | | | 18 -84 24 | | | | | | 6.43 | | 0.0016 | | | | |
|  |  | | Frontal_Mid_L | | | | -30 54 24 | | | | | | 6.36 | | 0.0016 | | | | |
|  |  | | Frontal_Sup_Orb_R | | | | 30 60 -6 | | | | | | 6.35 | | 0.0016 | | | | |
|  |  | | Rectus_R | | | | 12 48 -18 | | | | | | 6.25 | | 0.0022 | | | | |
|  |  | | Temporal_Mid_R | | | | 54 -42 -12 | | | | | | 6.18 | | 0.0026 | | | | |
|  |  | | Insula_L | | | | -42 18 0 | | | | | | 6.16 | | 0.0026 | | | | |
|  |  | | Lingual_R | | | | 24 -90 -6 | | | | | | 6.16 | | 0.0026 | | | | |
|  |  | | Frontal_Inf_Tri_L | | | | -36 24 18 | | | | | | 6.08 | | 0.0028 | | | | |
|  |  | | Angular_L | | | | -54 -66 24 | | | | | | 6.08 | | 0.0028 | | | | |
|  | 0.75 N2O | | SupraMarginal_R | | | | 48 -36 36 | | | | | | 7.26 | | 0.0007 | | | | |
|  |  | | Parietal_Inf_R | | | | 54 -36 48 | | | | | | 7.20 | | 0.0008 | | | | |
|  |  | | Postcentral_R | | | | 36 -30 42 | | | | | | 6.80 | | 0.0012 | | | | |
|  |  | | Precuneus_R | | | | 18 -60 42 | | | | | | 6.70 | | 0.0014 | | | | |
|  |  | | Precentral_R | | | | 30 -24 60 | | | | | | 6.65 | | 0.0014 | | | | |
|  |  | | Occipital_Sup_R | | | | 24 -66 36 | | | | | | 6.63 | | 0.0014 | | | | |
|  |  | | Postcentral_L | | | | -60 -18 30 | | | | | | 6.48 | | 0.0016 | | | | |
|  |  | | Parietal_Sup_R | | | | 24 -60 48 | | | | | | 6.41 | | 0.0018 | | | | |
|  |  | | Cuneus_R | | | | 18 -72 36 | | | | | | 6.34 | | 0.0018 | | | | |
|  |  | | Occipital_Mid_L | | | | -24 -78 36 | | | | | | 6.27 | | 0.0018 | | | | |
|  |  | | Precentral_L | | | | -48 0 36 | | | | | | 6.21 | | 0.0018 | | | | |
|  |  | | Cuneus_L | | | | -18 -78 36 | | | | | | 6.07 | | 0.0026 | | | | |
|  |  | | Precuneus_L | | | | 0 -48 54 | | | | | | 5.99 | | 0.0026 | | | | |
|  |  | | Angular_R | | | | 42 -54 36 | | | | | | 5.97 | | 0.0026 | | | | |
|  |  | | Occipital_Sup_L | | | | -18 -78 30 | | | | | | 5.78 | | 0.0040 | | | | |
|  | **Theta**  **MAC-awake Level** | | **Region of Interest** | | | | | **Voxel Coordinate** | | | **t-value** | | | | | | | | **p-value** |
|  | 1.30 Xenon | | Parietal_Sup_L | | | | | -30 -66 48 | | | 8.52 | | | | | | | | 0.0003 |
|  |  | | Precuneus_L | | | | | -6 -78 54 | | | 8.40 | | | | | | | | 0.0004 |
|  |  | | SupraMarginal_R | | | | | 60 -30 24 | | | 8.32 | | | | | | | | 0.0004 |
|  |  | | Paracentral_Lobule_R | | | | | 6 -42 60 | | | 8.22 | | | | | | | | 0.0004 |
|  |  | | Frontal_Sup_R | | | | | 24 24 60 | | | 8.20 | | | | | | | | 0.0004 |
|  |  | | Parietal_Sup_R | | | | | 18 -60 54 | | | 8.15 | | | | | | | | 0.0004 |
|  |  | | Precuneus_R | | | | | 12 -66 42 | | | 8.09 | | | | | | | | 0.0004 |
|  |  | | Postcentral_R | | | | | 12 -36 66 | | | 8.08 | | | | | | | | 0.0004 |
|  |  | | Cuneus_L | | | | | -6 -78 36 | | | 7.99 | | | | | | | | 0.0004 |
|  |  | | Occipital_Sup_L | | | | | -6 -84 48 | | | 7.81 | | | | | | | | 0.0006 |
|  |  | | Parietal_Inf_L | | | | | -30 -72 48 | | | 7.78 | | | | | | | | 0.0008 |
|  |  | | Precentral_R | | | | | 60 -6 42 | | | 7.76 | | | | | | | | 0.0008 |
|  |  | | Parietal_Inf_R | | | | | 42 -48 42 | | | 7.73 | | | | | | | | 0.0008 |
|  |  | | Frontal_Mid_R | | | | | 24 18 54 | | | 7.69 | | | | | | | | 0.0008 |
|  |  | | Angular_R | | | | | 42 -54 36 | | | 7.49 | | | | | | | | 0.0010 |
|  |  | | Cingulum_Mid_L | | | | | -6 -42 54 | | | 7.42 | | | | | | | | 0.0010 |
|  |  | | Temporal_Sup_R | | | | | 54 -30 18 | | | 7.38 | | | | | | | | 0.0010 |
|  |  | | Cuneus_R | | | | | 18 -66 30 | | | 7.27 | | | | | | | | 0.0010 |
|  |  | | Frontal_Mid_Orb_R | | | | | 36 48 -12 | | | 7.19 | | | | | | | | 0.0010 |
|  |  | | Supp_Motor_Area_R | | | | | 12 18 54 | | | 7.17 | | | | | | | | 0.0012 |
|  |  | | Frontal_Sup_Medial_R | | | | | 12 24 60 | | | 7.15 | | | | | | | | 0.0012 |
|  |  | | Paracentral_Lobule_L | | | | | -6 -30 54 | | | 7.02 | | | | | | | | 0.0014 |
|  |  | | Cingulum_Mid_R | | | | | 6 -12 48 | | | 6.82 | | | | | | | | 0.0016 |
|  |  | | Frontal_Med_Orb_R | | | | | 6 42 -6 | | | 6.76 | | | | | | | | 0.0016 |
|  |  | | Occipital_Sup_R | | | | | 30 -66 42 | | | 6.69 | | | | | | | | 0.0016 |
|  |  | | Supp_Motor_Area_L | | | | | 0 0 72 | | | 6.65 | | | | | | | | 0.0016 |
|  |  | | Temporal_Pole_Mid_R | | | | | 54 6 -24 | | | 6.65 | | | | | | | | 0.0016 |
|  |  | | Occipital_Mid_R | | | | | 30 -66 36 | | | 6.46 | | | | | | | | 0.0020 |
|  |  | | Frontal_Sup_Medial_L | | | | | 0 30 60 | | | 6.45 | | | | | | | | 0.0020 |
|  |  | | Frontal_Inf_Orb_R | | | | | 42 48 -12 | | | 6.44 | | | | | | | | 0.0020 |
|  |  | | Frontal_Sup_L | | | | | -12 0 72 | | | 6.38 | | | | | | | | 0.0022 |
|  |  | | Temporal_Pole_Sup_R | | | | | 48 12 -24 | | | 6.34 | | | | | | | | 0.0024 |
|  |  | | Frontal_Sup_Orb_R | | | | | 18 54 -12 | | | 6.29 | | | | | | | | 0.0024 |
|  |  | | Pallidum_R | | | | | 24 -6 6 | | | 6.21 | | | | | | | | 0.0024 |
|  |  | | Caudate_R | | | | | 18 -6 18 | | | 6.20 | | | | | | | | 0.0024 |
|  |  | | Rolandic_Oper_R | | | | | 48 -30 18 | | | 6.19 | | | | | | | | 0.0024 |
|  |  | | Postcentral_L | | | | | -18 -36 66 | | | 6.18 | | | | | | | | 0.0024 |
|  |  | | Frontal_Inf_Tri_R | | | | | 36 30 24 | | | 6.14 | | | | | | | | 0.0024 |
|  |  | | Occipital_Mid_L | | | | | -24 -78 36 | | | 6.06 | | | | | | | | 0.0028 |
|  |  | | Thalamus_R | | | | | 6 -18 12 | | | 6.05 | | | | | | | | 0.0028 |
|  |  | | Frontal_Inf_Oper_R | | | | | 48 18 36 | | | 5.99 | | | | | | | | 0.0030 |
|  |  | | Calcarine_L | | | | | 0 -96 12 | | | 5.95 | | | | | | | | 0.0032 |
|  |  | | Frontal_Med_Orb_L | | | | | 0 42 -12 | | | 5.85 | | | | | | | | 0.0036 |
|  |  | | Angular_L | | | | | -36 -66 42 | | | 5.76 | | | | | | | | 0.0036 |
|  |  | | Putamen_R | | | | | 24 0 12 | | | 5.73 | | | | | | | | 0.0036 |
|  |  | | Cingulum_Ant_R | | | | | 6 36 0 | | | 5.72 | | | | | | | | 0.0038 |
|  |  | | Calcarine_R | | | | | 12 -90 0 | | | 5.72 | | | | | | | | 0.0036 |
|  |  | | Temporal_Mid_R | | | | | 54 -66 18 | | | 5.70 | | | | | | | | 0.0040 |
|  | 0.75 Xenon | | Parietal_Inf_R | | | | | 54 -36 54 | | | 6.17 | | | | | | | | 0.0011 |
|  |  | | Precuneus_L | | | | | -6 -54 60 | | | 5.98 | | | | | | | | 0.0012 |
|  |  | | SupraMarginal_R | | | | | 48 -36 30 | | | 5.71 | | | | | | | | 0.0018 |
|  |  | | Postcentral_R | | | | | 48 -30 54 | | | 5.68 | | | | | | | | 0.0026 |
|  | 0.25 Xenon | | Parietal_Inf_L | | | | | -36 -60 48 | | | 6.14 | | | | | | | | 0.0019 |
|  | **Alpha**  **MAC-awake Level** | | **Region of Interest** | | | | | **Voxel Coordinate** | | | **t-value** | | | | | **p-value** | | | |
|  | 0.75 N2O | | Frontal_Sup_Medial_L | | | | | 0 66 0 | | | -9.92 | | | | | 0.0001 | | | |
|  |  | | Postcentral_L | | | | | -60 -18 30 | | | -9.78 | | | | | 0.0002 | | | |
|  |  | | Frontal_Mid_L | | | | | -42 54 6 | | | -9.70 | | | | | 0.0002 | | | |
|  |  | | Frontal_Inf_Tri_L | | | | | -48 36 18 | | | -9.60 | | | | | 0.0002 | | | |
|  |  | | Rectus_R | | | | | 6 36 -24 | | | -9.58 | | | | | 0.0002 | | | |
|  |  | | Cingulum_Ant_L | | | | | 0 36 30 | | | -9.58 | | | | | 0.0002 | | | |
|  |  | | SupraMarginal_L | | | | | -60 -24 24 | | | -9.50 | | | | | 0.0002 | | | |
|  |  | | Frontal_Sup_Orb_R | | | | | 12 18 -18 | | | -9.38 | | | | | 0.0002 | | | |
|  |  | | Cingulum_Mid_R | | | | | 6 36 36 | | | -9.21 | | | | | 0.0002 | | | |
|  |  | | Olfactory_R | | | | | 6 18 -18 | | | -9.14 | | | | | 0.0002 | | | |
|  |  | | Rectus_L | | | | | 0 24 -24 | | | -8.96 | | | | | 0.0002 | | | |
|  |  | | Frontal_Inf_Orb_R | | | | | 30 30 -24 | | | -8.83 | | | | | 0.0002 | | | |
|  |  | | Frontal_Sup_Orb_L | | | | | -12 36 -24 | | | -8.82 | | | | | 0.0002 | | | |
|  |  | | Putamen_L | | | | | -24 -6 12 | | | -8.72 | | | | | 0.0002 | | | |
|  |  | | Temporal_Sup_L | | | | | -54 -30 18 | | | -8.65 | | | | | 0.0002 | | | |
|  |  | | Frontal_Med_Orb_R | | | | | 6 24 -12 | | | -8.64 | | | | | 0.0002 | | | |
|  |  | | Frontal_Inf_Oper_R | | | | | 48 12 18 | | | -8.62 | | | | | 0.0002 | | | |
|  |  | | Frontal_Mid_Orb_R | | | | | 24 36 -18 | | | -8.58 | | | | | 0.0002 | | | |
|  |  | | Frontal_Sup_Medial_R | | | | | 6 42 36 | | | -8.56 | | | | | 0.0002 | | | |
|  |  | | Caudate_R | | | | | 12 12 -12 | | | -8.46 | | | | | 0.0002 | | | |
|  |  | | Frontal_Sup_L | | | | | -30 54 0 | | | -8.45 | | | | | 0.0002 | | | |
|  |  | | Frontal_Mid_R | | | | | 36 54 0 | | | -8.44 | | | | | 0.0002 | | | |
|  |  | | Rolandic_Oper_L | | | | | -48 -12 18 | | | -8.32 | | | | | 0.0002 | | | |
|  |  | | Frontal_Sup_R | | | | | 24 36 36 | | | -8.24 | | | | | 0.0002 | | | |
|  |  | | Precentral_L | | | | | -54 6 24 | | | -8.23 | | | | | 0.0002 | | | |
|  |  | | Insula_R | | | | | 36 18 -12 | | | -8.23 | | | | | 0.0002 | | | |
|  |  | | Frontal_Inf_Oper_L | | | | | -48 12 18 | | | -8.09 | | | | | 0.0002 | | | |
|  |  | | Insula_L | | | | | -36 -12 18 | | | -8.06 | | | | | 0.0002 | | | |
|  |  | | Temporal_Pole_Sup_L | | | | | -48 12 -6 | | | -8.03 | | | | | 0.0002 | | | |
|  |  | | Temporal_Inf_R | | | | | 54 -18 -30 | | | -7.75 | | | | | 0.0002 | | | |
|  |  | | Fusiform_L | | | | | -24 0 -42 | | | -7.70 | | | | | 0.0002 | | | |
|  |  | | Temporal_Pole_Sup_R | | | | | 30 24 -30 | | | -7.66 | | | | | 0.0002 | | | |
|  |  | | Putamen_R | | | | | 18 18 -6 | | | -7.65 | | | | | 0.0002 | | | |
|  |  | | ParaHippocampal_L | | | | | -24 -18 -24 | | | -7.57 | | | | | 0.0002 | | | |
|  |  | | Frontal_Inf_Orb_L | | | | | -18 24 -24 | | | -7.55 | | | | | 0.0002 | | | |
|  |  | | Temporal_Mid_L | | | | | -60 -6 -18 | | | -7.55 | | | | | 0.0002 | | | |
|  |  | | Hippocampus_L | | | | | -18 -24 -12 | | | -7.53 | | | | | 0.0002 | | | |
|  |  | | Frontal_Mid_Orb_L | | | | | -18 30 -18 | | | -7.45 | | | | | 0.0002 | | | |
|  |  | | Temporal_Pole_Mid_L | | | | | -42 18 -30 | | | -7.43 | | | | | 0.0002 | | | |
|  |  | | Olfactory_L | | | | | -6 18 -18 | | | -7.34 | | | | | 0.0002 | | | |
|  |  | | Parietal_Inf_L | | | | | -60 -36 42 | | | -7.33 | | | | | 0.0002 | | | |
|  |  | | Frontal_Inf_Tri_R | | | | | 42 12 24 | | | -7.32 | | | | | 0.0002 | | | |
|  |  | | Temporal_Mid_R | | | | | 54 -12 -18 | | | -7.30 | | | | | 0.0002 | | | |
|  |  | | Temporal_Inf_L | | | | | -60 -54 -6 | | | -7.14 | | | | | 0.0002 | | | |
|  |  | | Caudate_L | | | | | -18 0 18 | | | -7.01 | | | | | 0.0002 | | | |
|  |  | | Cingulum_Mid_L | | | | | -6 -24 42 | | | -6.96 | | | | | 0.0002 | | | |
|  |  | | Rolandic_Oper_R | | | | | 42 -12 18 | | | -6.94 | | | | | 0.0002 | | | |
|  |  | | Frontal_Med_Orb_L | | | | | -6 30 -12 | | | -6.90 | | | | | 0.0002 | | | |
|  |  | | Fusiform_R | | | | | 42 -72 -18 | | | -6.81 | | | | | 0.0002 | | | |
|  |  | | Precentral_R | | | | | 54 -6 48 | | | -6.79 | | | | | 0.0002 | | | |
|  |  | | Pallidum_R | | | | | 18 0 -6 | | | -6.73 | | | | | 0.0002 | | | |
|  |  | | Temporal_Pole_Mid_R | | | | | 42 18 -36 | | | -6.68 | | | | | 0.0002 | | | |
|  |  | | Temporal_Sup_R | | | | | 60 0 -12 | | | -6.64 | | | | | 0.0002 | | | |
|  |  | | Cingulum_Ant_R | | | | | 6 30 24 | | | -6.54 | | | | | 0.0004 | | | |
|  |  | | Supp_Motor_Area_R | | | | | 12 18 48 | | | -6.51 | | | | | 0.0004 | | | |
|  |  | | Postcentral_R | | | | | 60 -6 24 | | | -6.51 | | | | | 0.0004 | | | |
|  |  | | Supp_Motor_Area_L | | | | | -6 18 48 | | | -6.50 | | | | | 0.0004 | | | |
|  |  | | ParaHippocampal_R | | | | | 24 12 -30 | | | -6.47 | | | | | 0.0004 | | | |
|  |  | | Thalamus_L | | | | | -18 -12 6 | | | -6.31 | | | | | 0.0006 | | | |
|  |  | | Hippocampus_R | | | | | 18 6 12 | | | -5.97 | | | | | 0.0020 | | | |
|  |  | | Amygdala_L | | | | | -24 0 -24 | | | -5.93 | | | | | 0.0020 | | | |
|  |  | | Amygdala_R | | | | | 24 0 -12 | | | -5.93 | | | | | 0.0020 | | | |
|  |  | | Occipital_Inf_R | | | | | 42 -72 -12 | | | -5.87 | | | | | 0.0022 | | | |
|  |  | | Heschl_L | | | | | -36 -24 12 | | | -5.87 | | | | | 0.0022 | | | |
|  |  | | Lingual_R | | | | | 42 -78 -18 | | | -5.84 | | | | | 0.0022 | | | |
|  |  | | SupraMarginal_R | | | | | 54 -12 24 | | | -5.82 | | | | | 0.0022 | | | |
|  |  | | Thalamus_R | | | | | 6 -12 0 | | | -5.82 | | | | | 0.0022 | | | |
|  |  | | Pallidum_L | | | | | -24 -6 0 | | | -5.79 | | | | | 0.0022 | | | |
|  |  | | Heschl_R | | | | | 42 -18 12 | | | -5.79 | | | | | 0.0022 | | | |
|  |  | | Precuneus_R | | | | | 24 -48 0 | | | -5.61 | | | | | 0.0030 | | | |
|  |  | | Calcarine_L | | | | | 0 -84 -12 | | | -5.60 | | | | | 0.0030 | | | |
|  |  | | Paracentral_Lobule_L | | | | | -6 -24 54 | | | -5.58 | | | | | 0.0030 | | | |
|  |  | | Occipital_Inf_L | | | | | -48 -72 -18 | | | -5.52 | | | | | 0.0036 | | | |
|  |  | | Lingual_L | | | | | -12 -90 -18 | | | -5.51 | | | | | 0.0036 | | | |
|  |  | | Parietal_Inf_R | | | | | 48 -54 54 | | | -5.40 | | | | | 0.0042 | | | |
|  | 0.50 N2O | | Postcentral_L | | | | | -54 -18 30 | | | -7.23 | | | | | 0.0001 | | | |
|  |  | | Rectus_R | | | | | 6 36 -24 | | | -7.12 | | | | | 0.0002 | | | |
|  |  | | Rectus_L | | | | | -6 24 -24 | | | -6.98 | | | | | 0.0002 | | | |
|  |  | | Frontal_Sup_Medial_L | | | | | -12 54 0 | | | -6.82 | | | | | 0.0004 | | | |
|  |  | | Frontal_Inf_Tri_L | | | | | -48 36 18 | | | -6.81 | | | | | 0.0004 | | | |
|  |  | | Frontal_Sup_L | | | | | -18 60 6 | | | -6.76 | | | | | 0.0006 | | | |
|  |  | | Frontal_Inf_Tri_R | | | | | 54 30 18 | | | -6.65 | | | | | 0.0006 | | | |
|  |  | | Supp_Motor_Area_L | | | | | -6 18 48 | | | -6.63 | | | | | 0.0006 | | | |
|  |  | | Frontal_Inf_Oper_L | | | | | -54 12 18 | | | -6.61 | | | | | 0.0006 | | | |
|  |  | | SupraMarginal_L | | | | | -60 -24 30 | | | -6.59 | | | | | 0.0006 | | | |
|  |  | | Cingulum_Ant_L | | | | | -6 54 0 | | | -6.58 | | | | | 0.0006 | | | |
|  |  | | Frontal_Sup_Orb_R | | | | | 18 42 -24 | | | -6.57 | | | | | 0.0006 | | | |
|  |  | | Frontal_Inf_Orb_L | | | | | -42 36 -12 | | | -6.50 | | | | | 0.0006 | | | |
|  |  | | Frontal_Inf_Oper_R | | | | | 42 12 12 | | | -6.47 | | | | | 0.0006 | | | |
|  |  | | Lingual_L | | | | | -12 -90 -18 | | | -6.45 | | | | | 0.0006 | | | |
|  |  | | Frontal_Sup_Orb_L | | | | | -12 30 -24 | | | -6.44 | | | | | 0.0006 | | | |
|  |  | | Frontal_Mid_L | | | | | -42 54 6 | | | -6.41 | | | | | 0.0006 | | | |
|  |  | | Parietal_Inf_L | | | | | -60 -36 42 | | | -6.38 | | | | | 0.0006 | | | |
|  |  | | Olfactory_L | | | | | -6 18 -18 | | | -6.31 | | | | | 0.0010 | | | |
|  |  | | Frontal_Mid_R | | | | | 30 18 54 | | | -6.26 | | | | | 0.0012 | | | |
|  |  | | Caudate_R | | | | | 18 24 6 | | | -6.26 | | | | | 0.0012 | | | |
|  |  | | Frontal_Mid_Orb_L | | | | | -24 60 -12 | | | -6.24 | | | | | 0.0012 | | | |
|  |  | | Frontal_Inf_Orb_R | | | | | 48 30 -12 | | | -6.17 | | | | | 0.0012 | | | |
|  |  | | Frontal_Sup_Medial_R | | | | | 6 60 6 | | | -6.13 | | | | | 0.0012 | | | |
|  |  | | Temporal_Pole_Mid_L | | | | | -36 18 -36 | | | -6.13 | | | | | 0.0012 | | | |
|  |  | | Temporal_Inf_L | | | | | -42 6 -42 | | | -6.08 | | | | | 0.0016 | | | |
|  |  | | Frontal_Mid_Orb_R | | | | | 24 36 -18 | | | -6.05 | | | | | 0.0016 | | | |
|  |  | | Insula_R | | | | | 36 12 12 | | | -6.04 | | | | | 0.0016 | | | |
|  |  | | Insula_L | | | | | -36 -12 18 | | | -6.00 | | | | | 0.0016 | | | |
|  |  | | Rolandic_Oper_L | | | | | -42 -18 18 | | | -5.98 | | | | | 0.0016 | | | |
|  |  | | Rolandic_Oper_R | | | | | 42 6 12 | | | -5.98 | | | | | 0.0016 | | | |
|  |  | | Cingulum_Ant_R | | | | | 12 42 6 | | | -5.93 | | | | | 0.0016 | | | |
|  |  | | Frontal_Med_Orb_L | | | | | -6 60 -6 | | | -5.88 | | | | | 0.0016 | | | |
|  |  | | Fusiform_L | | | | | -24 0 -42 | | | -5.87 | | | | | 0.0016 | | | |
|  |  | | Temporal_Pole_Sup_L | | | | | -36 18 -30 | | | -5.87 | | | | | 0.0016 | | | |
|  |  | | Frontal_Sup_R | | | | | 18 42 42 | | | -5.78 | | | | | 0.0018 | | | |
|  |  | | SupraMarginal_R | | | | | 60 -36 24 | | | -5.67 | | | | | 0.0024 | | | |
|  |  | | Frontal_Med_Orb_R | | | | | 12 36 -12 | | | -5.65 | | | | | 0.0024 | | | |
|  |  | | Calcarine_L | | | | | 0 -84 -12 | | | -5.64 | | | | | 0.0024 | | | |
|  |  | | Cingulum_Mid_R | | | | | 12 18 42 | | | -5.63 | | | | | 0.0024 | | | |
|  |  | | Rectus_L | | | | | 0 24 -24 | | | -5.62 | | | | | 0.0033 | | | |
|  |  | | Fusiform_R | | | | | 42 -66 -18 | | | -5.60 | | | | | 0.0030 | | | |
|  |  | | Postcentral_R | | | | | 54 -18 30 | | | -5.52 | | | | | 0.0036 | | | |
|  |  | | Precentral_L | | | | | -42 -6 30 | | | -5.46 | | | | | 0.0040 | | | |
|  |  | | Caudate_L | | | | | -6 12 -12 | | | -5.43 | | | | | 0.0040 | | | |
|  |  | | ParaHippocampal_L | | | | | -18 0 -36 | | | -5.42 | | | | | 0.0042 | | | |
|  |  | | Supp_Motor_Area_R | | | | | 12 18 48 | | | -5.39 | | | | | 0.0046 | | | |
|  |  | | Temporal_Mid_R | | | | | 60 -36 -6 | | | -5.35 | | | | | 0.0050 | | | |
|  | 0.25 N2O | | Fusiform_R | | | | | 36 -72 -18 | | | -6.92 | | | | | 0.0003 | | | |
|  |  | | Frontal_Mid_R | | | | | 42 54 12 | | | -6.51 | | | | | 0.0006 | | | |
|  |  | | Frontal_Sup_L | | | | | -18 66 12 | | | -6.15 | | | | | 0.0006 | | | |
|  |  | | Frontal_Sup_Medial_L | | | | | -12 66 12 | | | -5.94 | | | | | 0.0008 | | | |
|  |  | | Rectus_L | | | | | 0 36 -24 | | | -5.84 | | | | | 0.0008 | | | |
|  |  | | Calcarine_L | | | | | 0 -84 -12 | | | -5.78 | | | | | 0.0008 | | | |
|  |  | | Postcentral_L | | | | | -60 -18 30 | | | -5.76 | | | | | 0.0008 | | | |
|  |  | | Parietal_Inf_L | | | | | -60 -36 42 | | | -5.76 | | | | | 0.0008 | | | |
|  |  | | Frontal_Sup_R | | | | | 24 42 48 | | | -5.72 | | | | | 0.0008 | | | |
|  |  | | Frontal_Sup_Medial_R | | | | | 6 60 0 | | | -5.71 | | | | | 0.0010 | | | |
|  |  | | SupraMarginal_L | | | | | -60 -36 36 | | | -5.63 | | | | | 0.0016 | | | |
|  |  | | Frontal_Mid_L | | | | | -30 36 48 | | | -5.62 | | | | | 0.0016 | | | |
|  |  | | Frontal_Sup_Orb_R | | | | | 18 36 -24 | | | -5.58 | | | | | 0.0018 | | | |
|  |  | | Frontal_Med_Orb_L | | | | | 0 36 -12 | | | -5.57 | | | | | 0.0018 | | | |
|  |  | | Frontal_Inf_Tri_L | | | | | -42 24 24 | | | -5.49 | | | | | 0.0020 | | | |
|  |  | | Frontal_Sup_Orb_L | | | | | -12 24 -24 | | | -5.45 | | | | | 0.0024 | | | |
|  |  | | Frontal_Inf_Oper_R | | | | | 54 12 6 | | | -5.41 | | | | | 0.0026 | | | |
|  |  | | Frontal_Inf_Orb_R | | | | | 24 24 -18 | | | -5.41 | | | | | 0.0026 | | | |
|  |  | | Frontal_Mid_Orb_L | | | | | -36 54 -12 | | | -5.38 | | | | | 0.0028 | | | |
|  |  | | Cingulum_Ant_L | | | | | -6 42 0 | | | -5.38 | | | | | 0.0028 | | | |
|  |  | | Frontal_Med_Orb_R | | | | | 12 60 -12 | | | -5.37 | | | | | 0.0028 | | | |
|  |  | | Temporal_Mid_L | | | | | -60 -12 -24 | | | -5.34 | | | | | 0.0030 | | | |
|  |  | | Temporal_Mid_R | | | | | 60 -12 -24 | | | -5.33 | | | | | 0.0030 | | | |
|  |  | | Rectus_R | | | | | 6 42 -24 | | | -5.32 | | | | | 0.0032 | | | |
|  |  | | Supp_Motor_Area_R | | | | | 6 -12 60 | | | -5.31 | | | | | 0.0032 | | | |
|  |  | | Frontal_Mid_Orb_R | | | | | 24 36 -18 | | | -5.27 | | | | | 0.0040 | | | |
|  |  | | Supp_Motor_Area_L | | | | | 0 12 48 | | | -5.23 | | | | | 0.0042 | | | |
|  |  | | Cingulum_Ant_R | | | | | 12 42 0 | | | -5.23 | | | | | 0.0044 | | | |
|  |  | | Temporal_Pole_Mid_L | | | | | -36 18 -36 | | | -5.23 | | | | | 0.0042 | | | |
|  |  | | Temporal_Inf_R | | | | | 54 -18 -30 | | | -5.23 | | | | | 0.0042 | | | |
|  |  | | Olfactory_L | | | | | -6 24 -12 | | | -5.20 | | | | | 0.0050 | | | |
| **Beta**  **MAC-awake Level** | | **Region of Interest** | | **Voxel Coordinate** | | **t-value** | | | | **p-value** | | | |  |  |  |  |  |  |
| 1.30 Xenon | | Postcentral_R | | 48 -30 54 | | 7.51 | | | | 0.0005 | | | |  |  |  |  |  |  |
|  | | SupraMarginal_R | | 60 -48 36 | | 7.17 | | | | 0.0008 | | | |  |  |  |  |  |  |
|  | | Parietal_Inf_R | | 48 -36 54 | | 6.78 | | | | 0.0008 | | | |  |  |  |  |  |  |
|  | | Parietal_Sup_L | | -30 -66 48 | | 6.77 | | | | 0.0008 | | | |  |  |  |  |  |  |
|  | | Parietal_Inf_L | | -60 -48 36 | | 6.62 | | | | 0.0012 | | | |  |  |  |  |  |  |
|  | | Cuneus_L | | -12 -84 30 | | 6.50 | | | | 0.0014 | | | |  |  |  |  |  |  |
|  | | Cuneus_R | | 12 -84 36 | | 6.47 | | | | 0.0014 | | | |  |  |  |  |  |  |
|  | | Parietal_Sup_R | | 18 -54 60 | | 6.47 | | | | 0.0014 | | | |  |  |  |  |  |  |
|  | | Angular_L | | -36 -60 42 | | 6.35 | | | | 0.0014 | | | |  |  |  |  |  |  |
|  | | Occipital_Sup_R | | 18 -84 36 | | 6.34 | | | | 0.0014 | | | |  |  |  |  |  |  |
|  | | Angular_R | | 54 -54 36 | | 6.33 | | | | 0.0014 | | | |  |  |  |  |  |  |
|  | | Occipital_Sup_L | | -18 -78 30 | | 6.24 | | | | 0.0014 | | | |  |  |  |  |  |  |
|  | | Temporal_Inf_L | | -54 -60 -24 | | 6.22 | | | | 0.0014 | | | |  |  |  |  |  |  |
|  | | Frontal_Sup_R | | 24 18 60 | | 5.98 | | | | 0.0028 | | | |  |  |  |  |  |  |
|  | | Frontal_Mid_R | | 36 24 30 | | 5.89 | | | | 0.0030 | | | |  |  |  |  |  |  |
|  | | Occipital_Mid_L | | -24 -78 36 | | 5.87 | | | | 0.0034 | | | |  |  |  |  |  |  |
| 0.75 Xenon | | Angular_R | | 42 -54 36 | | 5.35 | | | | 0.0041 | | | |  |  |  |  |  |  |
| 0.25 Xenon | | Parietal_Inf_L | | -36 -60 48 | | 5.55 | | | | 0.0035 | | | |  |  |  |  |  |  |
| 0.75 N2O | | Postcentral_R | | 42 -30 42 | | 6.06 | | | | 0.0023 | | | |  |  |  |  |  |  |
|  | | Precentral_R | | 42 -18 54 | | 5.89 | | | | 0.0036 | | | |  |  |  |  |  |  |
|  | | Angular_L | | -48 -66 42 | | 5.77 | | | | 0.0040 | | | |  |  |  |  |  |  |
|  | | Precuneus_L | | -6 -60 48 | | 5.67 | | | | 0.0046 | | | |  |  |  |  |  |  |
| 0.50 N2O | | Postcentral_R | | 42 -30 54 | | 5.63 | | | | 0.0049 | | | |  |  |  |  |  |  |
| **Low gamma**  **MAC-awake Level** | | **Region of Interest** | | | **Voxel Coordinate** | | | | **t-value** | | | **p-value** | | | | |  |  |  |
| 0.75 N2O | | Occipital_Sup_L | | | -12 -90 30 | | | | 9.58 | | | 0.0001 | | | | |  |  |  |
|  | | Occipital_Mid_R | | | 30 -90 18 | | | | 9.21 | | | 0.0002 | | | | |  |  |  |
|  | | Cuneus_L | | | -6 -90 30 | | | | 9.02 | | | 0.0002 | | | | |  |  |  |
|  | | Calcarine_R | | | 18 -90 6 | | | | 8.98 | | | 0.0002 | | | | |  |  |  |
|  | | Cuneus_R | | | 18 -96 12 | | | | 8.89 | | | 0.0002 | | | | |  |  |  |
|  | | Occipital_Sup_R | | | 24 -84 12 | | | | 8.84 | | | 0.0002 | | | | |  |  |  |
|  | | Calcarine_L | | | -18 -102 -6 | | | | 8.54 | | | 0.0002 | | | | |  |  |  |
|  | | Occipital_Inf_L | | | -24 -96 -6 | | | | 8.50 | | | 0.0002 | | | | |  |  |  |
|  | | Occipital_Mid_L | | | -30 -84 36 | | | | 8.42 | | | 0.0002 | | | | |  |  |  |
|  | | Postcentral_R | | | 30 -36 66 | | | | 8.33 | | | 0.0002 | | | | |  |  |  |
|  | | Precuneus_R | | | 6 -72 60 | | | | 8.25 | | | 0.0002 | | | | |  |  |  |
|  | | Angular_L | | | -42 -60 36 | | | | 8.12 | | | 0.0002 | | | | |  |  |  |
|  | | Parietal_Sup_L | | | -18 -72 48 | | | | 7.98 | | | 0.0002 | | | | |  |  |  |
|  | | Lingual_L | | | -24 -90 -18 | | | | 7.93 | | | 0.0002 | | | | |  |  |  |
|  | | Lingual_R | | | 18 -96 -12 | | | | 7.77 | | | 0.0002 | | | | |  |  |  |
|  | | Occipital_Inf_R | | | 30 -96 -6 | | | | 7.55 | | | 0.0002 | | | | |  |  |  |
|  | | Parietal_Inf_L | | | -48 -60 42 | | | | 7.55 | | | 0.0002 | | | | |  |  |  |
|  | | Frontal_Sup_R | | | 24 0 66 | | | | 7.51 | | | 0.0002 | | | | |  |  |  |
|  | | Precuneus_L | | | 0 -66 60 | | | | 7.51 | | | 0.0002 | | | | |  |  |  |
|  | | Supp_Motor_Area_R | | | 12 6 72 | | | | 7.42 | | | 0.0006 | | | | |  |  |  |
|  | | Frontal_Mid_L | | | -24 18 60 | | | | 7.39 | | | 0.0006 | | | | |  |  |  |
|  | | Frontal_Sup_Medial_L | | | -12 24 60 | | | | 7.38 | | | 0.0006 | | | | |  |  |  |
|  | | Frontal_Sup_L | | | -18 18 60 | | | | 7.37 | | | 0.0006 | | | | |  |  |  |
|  | | Angular_R | | | 36 -72 42 | | | | 7.31 | | | 0.0006 | | | | |  |  |  |
|  | | Parietal_Sup_R | | | 36 -48 60 | | | | 7.25 | | | 0.0006 | | | | |  |  |  |
|  | | Supp_Motor_Area_L | | | -12 18 60 | | | | 7.19 | | | 0.0006 | | | | |  |  |  |
|  | | Cingulum_Mid_L | | | 0 -12 36 | | | | 7.06 | | | 0.0006 | | | | |  |  |  |
|  | | Precentral_R | | | 36 -24 66 | | | | 7.05 | | | 0.0006 | | | | |  |  |  |
|  | | Fusiform_R | | | 30 -84 -6 | | | | 6.94 | | | 0.0006 | | | | |  |  |  |
|  | | Temporal_Mid_R | | | 54 -66 18 | | | | 6.69 | | | 0.0006 | | | | |  |  |  |
|  | | Cingulum_Mid_R | | | 0 -12 30 | | | | 6.54 | | | 0.0008 | | | | |  |  |  |
|  | | Frontal_Mid_R | | | 24 54 30 | | | | 6.53 | | | 0.0008 | | | | |  |  |  |
|  | | Frontal_Inf_Orb_L | | | -30 42 -18 | | | | 6.52 | | | 0.0010 | | | | |  |  |  |
|  | | Paracentral_Lobule_L | | | -12 -18 72 | | | | 6.45 | | | 0.0010 | | | | |  |  |  |
|  | | Frontal_Sup_Medial_R | | | 12 24 60 | | | | 6.41 | | | 0.0014 | | | | |  |  |  |
|  | | Temporal_Mid_L | | | -48 -66 18 | | | | 6.39 | | | 0.0014 | | | | |  |  |  |
|  | | Precentral_L | | | -18 -12 72 | | | | 6.34 | | | 0.0014 | | | | |  |  |  |
|  | | Parietal_Inf_R | | | 36 -48 54 | | | | 6.32 | | | 0.0014 | | | | |  |  |  |
|  | | Frontal_Mid_Orb_L | | | -30 42 -12 | | | | 6.24 | | | 0.0018 | | | | |  |  |  |
|  | | Frontal_Med_Orb_L | | | -6 42 -12 | | | | 6.11 | | | 0.0024 | | | | |  |  |  |
|  | | Postcentral_L | | | -24 -30 72 | | | | 5.97 | | | 0.0030 | | | | |  |  |  |
|  | | Cingulum_Ant_L | | | -6 24 30 | | | | 5.79 | | | 0.0044 | | | | |  |  |  |
|  | | SupraMarginal_L | | | -60 -18 42 | | | | 5.76 | | | 0.0044 | | | | |  |  |  |
|  | | Frontal_Inf_Tri_L | | | -48 24 12 | | | | 5.73 | | | 0.0048 | | | | |  |  |  |
| 0.50 N2O | | Cuneus_L | | | -6 -90 30 | | | | 5.50 | | | 0.0043 | | | | |  |  |  |
| 1.30 Xenon | | Occipital_Inf_R | | | 24 -96 -6 | | | | 7.01 | | | 0.0007 | | | | |  |  |  |
|  | | Frontal_Sup_Medial_R | | | 6 24 60 | | | | 6.87 | | | 0.0012 | | | | |  |  |  |
|  | | Supp_Motor_Area_L | | | 0 24 66 | | | | 6.86 | | | 0.0012 | | | | |  |  |  |
|  | | Lingual_R | | | 24 -90 -6 | | | | 6.80 | | | 0.0014 | | | | |  |  |  |
|  | | Calcarine_R | | | 18 -96 -6 | | | | 6.68 | | | 0.0018 | | | | |  |  |  |
|  | | Angular_L | | | -36 -66 42 | | | | 6.68 | | | 0.0018 | | | | |  |  |  |
|  | | Cuneus_L | | | -12 -84 36 | | | | 6.61 | | | 0.0024 | | | | |  |  |  |
|  | | Occipital_Sup_L | | | -18 -84 36 | | | | 6.61 | | | 0.0024 | | | | |  |  |  |
|  | | Frontal_Sup_R | | | 18 18 66 | | | | 6.60 | | | 0.0026 | | | | |  |  |  |
|  | | Supp_Motor_Area_R | | | 6 24 54 | | | | 6.59 | | | 0.0026 | | | | |  |  |  |
|  | | Frontal_Sup_Medial_L | | | 0 30 60 | | | | 6.59 | | | 0.0026 | | | | |  |  |  |
|  | | Occipital_Mid_L | | | -24 -78 36 | | | | 6.59 | | | 0.0026 | | | | |  |  |  |
|  | | Parietal_Inf_L | | | -54 -48 36 | | | | 6.51 | | | 0.0026 | | | | |  |  |  |
|  | | Parietal_Sup_R | | | 18 -54 60 | | | | 6.40 | | | 0.0026 | | | | |  |  |  |
|  | | SupraMarginal_R | | | 60 -48 36 | | | | 6.37 | | | 0.0030 | | | | |  |  |  |
|  | | Precentral_R | | | 60 -6 42 | | | | 6.31 | | | 0.0034 | | | | |  |  |  |
|  | | Parietal_Inf_R | | | 54 -54 42 | | | | 6.31 | | | 0.0034 | | | | |  |  |  |
|  | | Cuneus_R | | | 12 -84 36 | | | | 6.29 | | | 0.0036 | | | | |  |  |  |
|  | | Occipital_Sup_R | | | 18 -84 36 | | | | 6.29 | | | 0.0036 | | | | |  |  |  |
|  | | Occipital_Inf_L | | | -18 -96 -6 | | | | 6.25 | | | 0.0040 | | | | |  |  |  |
|  | | Fusiform_R | | | 48 -72 -18 | | | | 6.23 | | | 0.0040 | | | | |  |  |  |
| 0.75 Xenon | | Precentral_R | | | 60 -6 42 | | | | 6.19 | | | 0.0003 | | | | |  |  |  |
|  | | Postcentral_R | | | 60 -6 36 | | | | 6.05 | | | 0.0010 | | | | |  |  |  |
|  | | Temporal_Mid_R | | | 42 -66 0 | | | | 5.53 | | | 0.0042 | | | | |  |  |  |
| **High Gamma**  **MAC-awake Level** | | **Region of Interest** | | | **Voxel Coordinate** | | | | **t-value** | | | **p-value** | | | | | |  |  |
| 0.75 N2O | | Frontal_Mid_R | | | 24 54 30 | | | | 7.18 | | | 1.00E-04 | | | | | |  |  |
|  | | Frontal_Sup_Medial_L | | | -6 30 60 | | | | 6.94 | | | 0.0002 | | | | | |  |  |
|  | | Precuneus_R | | | 6 -78 48 | | | | 6.92 | | | 0.0002 | | | | | |  |  |
|  | | Frontal_Sup_R | | | 24 54 36 | | | | 6.83 | | | 0.0002 | | | | | |  |  |
|  | | Cingulum_Mid_L | | | 0 0 42 | | | | 6.81 | | | 0.0002 | | | | | |  |  |
|  | | Supp_Motor_Area_L | | | -6 0 48 | | | | 6.64 | | | 0.0004 | | | | | |  |  |
|  | | Frontal_Sup_L | | | -18 24 60 | | | | 6.48 | | | 0.0004 | | | | | |  |  |
|  | | Occipital_Mid_R | | | 36 -84 24 | | | | 6.45 | | | 0.0006 | | | | | |  |  |
|  | | Frontal_Mid_L | | | -24 18 60 | | | | 6.42 | | | 0.0006 | | | | | |  |  |
|  | | Frontal_Sup_Medial_R | | | 12 60 30 | | | | 6.41 | | | 0.0006 | | | | | |  |  |
|  | | Precuneus_L | | | -12 -66 60 | | | | 6.27 | | | 0.0012 | | | | | |  |  |
|  | | Cingulum_Mid_R | | | 6 0 42 | | | | 6.15 | | | 0.0012 | | | | | |  |  |
|  | | Parietal_Sup_L | | | -18 -66 60 | | | | 6.14 | | | 0.0012 | | | | | |  |  |
|  | | Supp_Motor_Area_R | | | 12 6 66 | | | | 6.08 | | | 0.0014 | | | | | |  |  |
|  | | Cuneus_R | | | 18 -90 12 | | | | 6.05 | | | 0.0014 | | | | | |  |  |
|  | | Occipital_Sup_L | | | -18 -90 30 | | | | 6.02 | | | 0.0014 | | | | | |  |  |
|  | | Angular_R | | | 48 -72 30 | | | | 5.95 | | | 0.0014 | | | | | |  |  |
|  | | Calcarine_R | | | 18 -90 6 | | | | 5.91 | | | 0.0016 | | | | | |  |  |
|  | | Parietal_Sup_R | | | 12 -66 66 | | | | 5.86 | | | 0.0018 | | | | | |  |  |
|  | | Occipital_Mid_L | | | -30 -96 12 | | | | 5.74 | | | 0.002999 | | | | | |  |  |
|  | | Cuneus_L | | | -12 -78 18 | | | | 5.73 | | | 0.002999 | | | | | |  |  |
|  | | Occipital_Sup_R | | | 24 -90 18 | | | | 5.73 | | | 0.002999 | | | | | |  |  |
|  | | Angular_L | | | -48 -66 42 | | | | 5.61 | | | 0.003199 | | | | | |  |  |
|  | | Cingulum_Ant_L | | | 0 0 30 | | | | 5.51 | | | 0.003599 | | | | | |  |  |
|  | | Calcarine_L | | | -6 -72 18 | | | | 5.5 | | | 0.003599 | | | | | |  |  |
|  | | Precentral_R | | | 36 -24 66 | | | | 5.47 | | | 0.003599 | | | | | |  |  |
|  | | Parietal_Inf_L | | | -48 -60 48 | | | | 5.45 | | | 0.004199 | | | | | |  |  |
| 1.30 Xenon | | Rolandic_Oper_R | | | 48 0 6 | | | | 6.31 | | | 0.0005 | | | | | |  |  |

*Supp. Table 2B. Electroencephalographic sources most significantly altered by Xe and N_2_O administration.* Significantly (p<0.005 for N_2_O; p<0.004 for Xe) changed regions of interest against post-antiemetic baseline for each frequency band. Voxel coordinates are in AAL atlas coordinate system along with associated labels^43^. [delta (1-4 Hz), theta (4-8 Hz), alpha (8-13 Hz), low gamma (30-49 Hz), high gamma (51-99 Hz)].

| **Delta**  **MAC-awake Level** | | **Region of Interest** | | **Voxel Coordinate** | | | **t-value** | | | **p-value** | | |
| --- | --- | --- | --- | --- | --- | --- | --- | --- | --- | --- | --- | --- |
|  | 1.30 Xenon | Supp_Motor_Area_R | | 6 0 72 | | | 9.66 | | | 0.0007 | | |
|  |  | Supp_Motor_Area_L | | 0 0 72 | | | 9.58 | | | 0.0008 | | |
|  |  | Postcentral_L | | -48 -36 60 | | | 8.89 | | | 0.0008 | | |
|  |  | Cingulum_Mid_R | | 0 -12 30 | | | 8.79 | | | 0.0008 | | |
|  |  | Parietal_Inf_L | | -42 -48 54 | | | 8.76 | | | 0.0008 | | |
|  |  | Frontal_Sup_R | | 18 -12 72 | | | 8.67 | | | 0.0008 | | |
|  |  | Cingulum_Mid_L | | 0 -12 36 | | | 8.66 | | | 0.001 | | |
|  |  | Cingulum_Post_L | | -6 -36 30 | | | 8.64 | | | 0.001 | | |
|  |  | Precentral_R | | 18 -18 72 | | | 8.59 | | | 0.001 | | |
|  |  | Cuneus_L | | 0 -72 24 | | | 8.51 | | | 0.001 | | |
|  |  | Precuneus_L | | -12 -60 30 | | | 8.51 | | | 0.001 | | |
|  |  | Cuneus_R | | 6 -72 24 | | | 8.44 | | | 0.001 | | |
|  |  | Frontal_Mid_R | | 36 60 0 | | | 8.41 | | | 0.001 | | |
|  |  | Cingulum_Ant_L | | -6 0 30 | | | 8.37 | | | 0.001 | | |
|  |  | Precuneus_R | | 6 -66 24 | | | 8.30 | | | 0.001 | | |
|  |  | Caudate_L | | -18 -18 24 | | | 8.26 | | | 0.001 | | |
|  |  | Calcarine_R | | 6 -72 18 | | | 8.25 | | | 0.001 | | |
|  |  | Thalamus_L | | -6 -18 18 | | | 8.22 | | | 0.001 | | |
|  |  | Calcarine_L | | 0 -72 18 | | | 8.19 | | | 0.001 | | |
|  |  | Paracentral_Lobule_L | | 0 -24 54 | | | 8.18 | | | 0.001 | | |
|  |  | Postcentral_R | | 24 -36 72 | | | 8.09 | | | 0.001 | | |
|  |  | Paracentral_Lobule_R | | 12 -24 72 | | | 8.09 | | | 0.001 | | |
|  |  | Frontal_Sup_L | | -12 -6 72 | | | 7.93 | | | 0.001 | | |
|  |  | Parietal_Sup_R | | 18 -54 72 | | | 7.91 | | | 0.001 | | |
|  |  | Thalamus_R | | 12 -18 18 | | | 7.88 | | | 0.001 | | |
|  |  | Cingulum_Post_R | | 6 -54 30 | | | 7.80 | | | 0.001 | | |
|  |  | Angular_L | | -30 -54 36 | | | 7.73 | | | 0.001 | | |
|  |  | Caudate_R | | 18 -12 24 | | | 7.72 | | | 0.001 | | |
|  |  | Occipital_Sup_R | | 24 -72 18 | | | 7.68 | | | 0.001 | | |
|  |  | Temporal_Inf_R | | 36 0 -42 | | | 7.68 | | | 0.001 | | |
|  |  | Precentral_L | | -36 -30 60 | | | 7.67 | | | 0.001 | | |
|  |  | Parietal_Sup_L | | -30 -48 60 | | | 7.66 | | | 0.001 | | |
|  |  | SupraMarginal_L | | -60 -24 42 | | | 7.57 | | | 0.001 | | |
|  |  | Occipital_Mid_R | | 36 -78 12 | | | 7.47 | | | 0.001 | | |
|  |  | Occipital_Sup_L | | -18 -66 30 | | | 7.39 | | | 0.0012 | | |
|  |  | Lingual_R | | 6 -66 6 | | | 7.37 | | | 0.0012 | | |
|  |  | Occipital_Mid_L | | -24 -60 36 | | | 7.33 | | | 0.0012 | | |
|  |  | Temporal_Sup_R | | 66 -24 12 | | | 7.29 | | | 0.0012 | | |
|  |  | Lingual_L | | 0 -72 6 | | | 7.26 | | | 0.0012 | | |
|  |  | Frontal_Inf_Tri_R | | 48 48 0 | | | 7.25 | | | 0.0012 | | |
|  |  | Frontal_Mid_Orb_R | | 42 54 -6 | | | 7.19 | | | 0.0012 | | |
|  |  | SupraMarginal_R | | 66 -18 24 | | | 7.19 | | | 0.0012 | | |
|  |  | Frontal_Mid_L | | -42 54 6 | | | 7.11 | | | 0.0012 | | |
|  |  | Insula_L | | -36 -12 18 | | | 7.02 | | | 0.0014 | | |
|  |  | Putamen_L | | -24 -6 12 | | | 6.99 | | | 0.0014 | | |
|  |  | Fusiform_R | | 30 06 36 | | | 6.97 | | | 0.0014 | | |
|  |  | Parietal_Inf_R | | 60 -36 48 | | | 6.91 | | | 0.0014 | | |
|  |  | Occipital_Inf_L | | -48 -72 -18 | | | 6.83 | | | 0.0014 | | |
|  |  | Rolandic_Oper_L | | -36 -24 18 | | | 6.81 | | | 0.0014 | | |
|  |  | Rolandic_Oper_R | | 60 -18 18 | | | 6.78 | | | 0.0014 | | |
|  |  | Pallidum_L | | -18 0 6 | | | 6.53 | | | 0.0014 | | |
|  |  | Heschl_L | | -36 -24 12 | | | 6.51 | | | 0.0014 | | |
|  |  | Occipital_Inf_R | | 30 -96 -6 | | | 6.49 | | | 0.0014 | | |
|  |  | Temporal_Sup_L | | -42 -42 18 | | | 6.48 | | | 0.0014 | | |
|  |  | Fusiform_L | | -42 -72 -18 | | | 6.43 | | | 0.0014 | | |
|  |  | Insula_R | | 30 -18 18 | | | 6.42 | | | 0.0014 | | |
|  |  | Temporal_Mid_R | | 54 -36 -12 | | | 6.41 | | | 0.0014 | | |
|  |  | ParaHippocampal_R | | 30 06 30 | | | 6.40 | | | 0.0014 | | |
|  |  | Frontal_Inf_Tri_L | | -48 42 12 | | | 6.39 | | | 0.0014 | | |
|  |  | Temporal_Pole_Mid_R | | 36 6 -36 | | | 6.39 | | | 0.0014 | | |
|  |  | Hippocampus_R | | 36 -30 -12 | | | 6.28 | | | 0.0014 | | |
|  |  | Hippocampus_L | | -18 -36 6 | | | 6.22 | | | 0.0014 | | |
|  |  | Temporal_Mid_L | | -48 -48 18 | | | 6.20 | | | 0.0014 | | |
|  |  | Temporal_Pole_Sup_R | | 42 18 -30 | | | 6.17 | | | 0.0016 | | |
|  |  | Frontal_Sup_Medial_L | | -6 18 42 | | | 6.14 | | | 0.0016 | | |
|  |  | Temporal_Inf_L | | -54 -66 -12 | | | 6.12 | | | 0.0016 | | |
|  |  | Frontal_Mid_Orb_L | | -42 48 -6 | | | 6.10 | | | 0.0016 | | |
|  |  | Angular_R | | 30 -54 42 | | | 6.08 | | | 0.0016 | | |
|  |  | ParaHippocampal_L | | -18 -36 -12 | | | 6.02 | | | 0.0018 | | |
|  |  | Amygdala_R | | 30 0 -24 | | | 6.02 | | | 0.0018 | | |
|  |  | Putamen_R | | 30 -18 0 | | | 5.80 | | | 0.0024 | | |
|  |  | Frontal_Inf_Orb_R | | 36 24 -24 | | | 5.78 | | | 0.0024 | | |
|  |  | Frontal_Sup_Medial_R | | 12 60 30 | | | 5.77 | | | 0.0024 | | |
|  |  | Frontal_Inf_Oper_L | | -36 6 24 | | | 5.74 | | | 0.0026 | | |
|  |  | Frontal_Sup_Orb_R | | 30 60 -6 | | | 5.73 | | | 0.0026 | | |
|  |  | Pallidum_R | | 30 12 06 | | | 5.71 | | | 0.0026 | | |
|  |  | Heschl_R | | 42 -24 12 | | | 5.61 | | | 0.0026 | | |
|  |  | Frontal_Inf_Oper_R | | 30 6 30 | | | 5.60 | | | 0.0026 | | |
|  |  | Frontal_Inf_Orb_L | | -42 42 -6 | | | 5.48 | | | 0.002799 | | |
|  |  | Olfactory_L | | 0 12 -6 | | | 5.46 | | | 0.002799 | | |
|  |  | Rectus_L | | -12 24 -12 | | | 5.40 | | | 0.003399 | | |
|  |  | Olfactory_R | | 24 12 -18 | | | 5.38 | | | 0.003599 | | |
|  | 0.75 Xenon | Parietal_Sup_L | | -24 -78 48 | | | 5.36 | | | 0.002899 | | |
|  |  | Precuneus_L | | -12 -66 48 | | | 5.25 | | | 0.002999 | | |
|  |  | Parietal_Inf_L | | -30 -78 48 | | | 4.99 | | | 0.004399 | | |
|  |  | Parietal_Inf_R | | 48 -54 54 | | | 4.26 | | | 0.0025 | | |
|  |  | Parietal_Inf_L | | -60 -30 42 | | | 4.15 | | | 0.003199 | | |
|  |  | SupraMarginal_L | | -60 -30 36 | | | 4.15 | | | 0.003199 | | |
|  |  | SupraMarginal_R | | 60 -48 24 | | | 4.13 | | | 0.003199 | | |
|  |  | Temporal_Mid_R | | 60 -30 -12 | | | 4.03 | | | 0.003599 | | |
|  |  | Parietal_Sup_L | | -24 -48 72 | | | 4.01 | | | 0.003599 | | |
|  |  | Angular_R | | 54 -48 30 | | | 4.01 | | | 0.003599 | | |
|  |  | Postcentral_R | | 42 -36 54 | | | 3.99 | | | 0.003799 | | |
|  |  | Postcentral_L | | -24 -36 72 | | | 3.97 | | | 0.003799 | | |
|  |  | Temporal_Sup_R | | 60 -42 18 | | | 3.92 | | | 0.003999 | | |
|  | **Theta**  **MAC-awake Level** | | **Region of Interest** | | **Voxel Coordinate** | | | **t-value** | | | **p-value** | |
|  | 1.30 Xenon | | Postcentral_R | | 30 -36 72 | | | 8.26 | | | 1E-04 | |
|  |  | | Parietal_Sup_L | | -24 -48 72 | | | 7.50 | | | 0.0002 | |
|  |  | | Paracentral_Lobule_L | | -12 -36 78 | | | 7.47 | | | 0.0002 | |
|  |  | | Occipital_Inf_L | | -48 -66 -18 | | | 7.33 | | | 0.0002 | |
|  |  | | Temporal_Inf_L | | -54 -66 -12 | | | 7.17 | | | 0.0002 | |
|  |  | | Fusiform_L | | -42 -66 -18 | | | 7.02 | | | 0.0002 | |
|  |  | | Supp_Motor_Area_R | | 12 6 72 | | | 6.97 | | | 0.0004 | |
|  |  | | Occipital_Mid_L | | -54 -72 0 | | | 6.85 | | | 0.0004 | |
|  |  | | Frontal_Sup_Medial_R | | 12 54 42 | | | 6.79 | | | 0.0004 | |
|  |  | | Precuneus_L | | -12 -42 78 | | | 6.73 | | | 0.0004 | |
|  |  | | Temporal_Mid_L | | -60 -60 0 | | | 6.70 | | | 0.0004 | |
|  |  | | Frontal_Sup_R | | 24 12 66 | | | 6.58 | | | 0.0004 | |
|  |  | | Precentral_R | | 30 -24 72 | | | 6.57 | | | 0.0004 | |
|  |  | | Precentral_L | | -24 -24 72 | | | 6.48 | | | 0.0004 | |
|  |  | | Temporal_Sup_L | | -60 -24 12 | | | 6.45 | | | 0.0004 | |
|  |  | | Paracentral_Lobule_R | | 6 -42 78 | | | 6.41 | | | 0.0004 | |
|  |  | | Postcentral_L | | -18 -30 72 | | | 6.40 | | | 0.0004 | |
|  |  | | Parietal_Inf_R | | 54 -42 54 | | | 6.30 | | | 0.0006 | |
|  |  | | Cuneus_R | | 18 -84 42 | | | 6.28 | | | 0.0006 | |
|  |  | | Supp_Motor_Area_L | | -12 -12 72 | | | 6.26 | | | 0.0012 | |
|  |  | | Occipital_Sup_R | | 24 -84 42 | | | 6.22 | | | 0.0012 | |
|  |  | | SupraMarginal_L | | -60 -24 18 | | | 6.22 | | | 0.0012 | |
|  |  | | Rolandic_Oper_L | | -48 -18 18 | | | 6.19 | | | 0.0012 | |
|  |  | | Insula_L | | -36 -18 18 | | | 5.99 | | | 0.0012 | |
|  |  | | Frontal_Sup_L | | -12 -6 72 | | | 5.93 | | | 0.0012 | |
|  |  | | Heschl_L | | -48 -12 6 | | | 5.89 | | | 0.0012 | |
|  |  | | Calcarine_R | | 12 -96 6 | | | 5.87 | | | 0.0012 | |
|  |  | | Parietal_Sup_R | | 24 -48 72 | | | 5.86 | | | 0.0012 | |
|  |  | | Frontal_Mid_Orb_L | | -42 54 -6 | | | 5.83 | | | 0.0012 | |
|  |  | | Precuneus_R | | 6 -60 72 | | | 5.79 | | | 0.0012 | |
|  |  | | Cuneus_L | | 0 -96 18 | | | 5.68 | | | 0.0014 | |
|  |  | | Occipital_Sup_L | | -6 -102 6 | | | 5.68 | | | 0.0014 | |
|  |  | | Angular_R | | 36 -78 42 | | | 5.67 | | | 0.0014 | |
|  |  | | Calcarine_L | | 0 -96 12 | | | 5.66 | | | 0.0014 | |
|  |  | | Occipital_Mid_R | | 30 -84 36 | | | 5.66 | | | 0.0014 | |
|  |  | | Lingual_L | | -30 -48 -6 | | | 5.62 | | | 0.0014 | |
|  |  | | Putamen_L | | -30 -18 6 | | | 5.49 | | | 0.0014 | |
|  |  | | Frontal_Mid_R | | 24 54 30 | | | 5.44 | | | 0.0014 | |
|  |  | | Temporal_Pole_Sup_L | | -54 6 -12 | | | 5.44 | | | 0.0014 | |
|  |  | | SupraMarginal_R | | 60 -30 48 | | | 5.43 | | | 0.0014 | |
|  |  | | ParaHippocampal_L | | -30 -42 -6 | | | 5.42 | | | 0.0014 | |
|  |  | | Frontal_Inf_Tri_L | | -48 18 0 | | | 5.39 | | | 0.0018 | |
|  |  | | Hippocampus_L | | -30 -36 0 | | | 5.38 | | | 0.002 | |
|  |  | | Frontal_Inf_Orb_L | | -48 24 -6 | | | 5.37 | | | 0.0026 | |
|  |  | | Frontal_Mid_L | | -42 48 18 | | | 5.34 | | | 0.0026 | |
|  |  | | Thalamus_L | | -18 -18 6 | | | 5.32 | | | 0.002799 | |
|  |  | | Frontal_Sup_Medial_L | | 0 66 6 | | | 5.28 | | | 0.002999 | |
|  |  | | Parietal_Inf_L | | -48 -24 36 | | | 5.24 | | | 0.003399 | |
|  |  | | Frontal_Inf_Oper_L | | -36 6 24 | | | 5.22 | | | 0.003399 | |
|  |  | | Cingulum_Mid_L | | -12 -36 54 | | | 5.21 | | | 0.003399 | |
|  |  | | Caudate_L | | -18 -18 24 | | | 5.20 | | | 0.003399 | |
|  |  | | Pallidum_L | | -24 -12 0 | | | 5.20 | | | 0.003399 | |
|  |  | | Frontal_Med_Orb_R | | 6 66 -6 | | | 5.13 | | | 0.003799 | |
|  |  | | Fusiform_R | | 42 -36 -24 | | | 5.09 | | | 0.003799 | |
|  |  | | Frontal_Med_Orb_L | | -6 66 -6 | | | 5.07 | | | 0.003799 | |
|  |  | | Cingulum_Ant_L | | -6 54 0 | | | 5.07 | | | 0.003799 | |
|  |  | | Cingulum_Post_L | | -6 -42 6 | | | 5.07 | | | 0.003799 | |
|  |  | | Frontal_Sup_Orb_L | | -12 60 -6 | | | 5.01 | | | 0.003999 | |
|  | **Alpha**  **MAC-awake Level** | | **Region of Interest** | | | **Voxel Coordinate** | | | **t-value** | | | **p-value** |
|  | 0.75 N2O | | Frontal_Sup_L | | | -12 6 72 | | | -4.65 | | | 0.0019 |
|  |  | | Precentral_L | | | -60 6 24 | | | -4.53 | | | 0.0022 |
|  |  | | Supp_Motor_Area_L | | | -6 6 72 | | | -4.47 | | | 0.0026 |
|  |  | | Frontal_Mid_L | | | -42 12 42 | | | -4.43 | | | 0.003199 |
|  |  | | Frontal_Inf_Tri_L | | | -48 24 24 | | | -4.42 | | | 0.003599 |
|  |  | | Temporal_Mid_L | | | -66 -36 -6 | | | -4.4 | | | 0.003799 |
|  |  | | Frontal_Inf_Oper_L | | | -60 6 18 | | | -4.37 | | | 0.003799 |
|  |  | | Postcentral_L | | | -60 0 18 | | | -4.37 | | | 0.003799 |
|  |  | | Frontal_Sup_R | | | 24 24 60 | | | -4.36 | | | 0.003799 |
|  | 0.50 Xenon | | Frontal_Sup_L | | | -18 -6 72 | | | -7.9 | | | 1.00E-04 |
|  |  | | Frontal_Sup_Medial_R | | | 12 48 48 | | | -7.78 | | | 0.0002 |
|  |  | | Frontal_Sup_R | | | 24 48 42 | | | -7.53 | | | 0.0004 |
|  |  | | Frontal_Mid_R | | | 36 42 36 | | | -7.5 | | | 0.0004 |
|  |  | | Frontal_Sup_Medial_L | | | 0 42 54 | | | -7.31 | | | 0.0004 |
|  |  | | Supp_Motor_Area_L | | | -6 6 72 | | | -6.88 | | | 0.0004 |
|  |  | | Temporal_Inf_L | | | -42 0 -42 | | | -6.76 | | | 0.0004 |
|  |  | | Frontal_Mid_L | | | -36 18 42 | | | -6.53 | | | 0.0004 |
|  |  | | Frontal_Inf_Tri_L | | | -48 24 30 | | | -6.5 | | | 0.0004 |
|  |  | | Fusiform_L | | | -36 -12 -42 | | | -6.4 | | | 0.0004 |
|  |  | | Precentral_L | | | -24 -12 72 | | | -6.3 | | | 0.0004 |
|  |  | | Paracentral_Lobule_L | | | -12 -24 78 | | | -6.2 | | | 0.0006 |
|  |  | | Supp_Motor_Area_R | | | 6 6 72 | | | -6.15 | | | 0.0006 |
|  |  | | Precentral_R | | | 42 -12 60 | | | -6.12 | | | 0.0006 |
|  |  | | Frontal_Inf_Tri_R | | | 48 30 30 | | | -6.12 | | | 0.0006 |
|  |  | | Temporal_Inf_R | | | 48 -48 -24 | | | -6.1 | | | 0.0008 |
|  |  | | Fusiform_R | | | 42 -42 -24 | | | -6.09 | | | 0.0008 |
|  |  | | Frontal_Inf_Oper_L | | | -54 12 24 | | | -6.02 | | | 0.0008 |
|  |  | | Parietal_Sup_L | | | -30 -60 66 | | | -5.98 | | | 0.0008 |
|  |  | | Temporal_Mid_R | | | 66 -42 6 | | | -5.98 | | | 0.0008 |
|  |  | | Temporal_Sup_R | | | 66 -36 12 | | | -5.95 | | | 0.0008 |
|  |  | | Frontal_Inf_Oper_R | | | 42 24 30 | | | -5.73 | | | 0.001 |
|  |  | | ParaHippocampal_L | | | -24 0 -36 | | | -5.71 | | | 0.001 |
|  |  | | Temporal_Pole_Mid_L | | | -30 6 -42 | | | -5.64 | | | 0.001 |
|  |  | | Temporal_Mid_L | | | -60 -6 -18 | | | -5.63 | | | 0.001 |
|  |  | | Cuneus_L | | | -6 -96 18 | | | -5.62 | | | 0.001 |
|  |  | | Postcentral_L | | | -60 0 18 | | | -5.62 | | | 0.001 |
|  |  | | Calcarine_L | | | -18 -102 -6 | | | -5.61 | | | 0.001 |
|  |  | | Occipital_Sup_L | | | -18 -90 24 | | | -5.61 | | | 0.001 |
|  |  | | Frontal_Med_Orb_L | | | -12 54 -6 | | | -5.58 | | | 0.0022 |
|  |  | | Rolandic_Oper_L | | | -42 0 18 | | | -5.5 | | | 0.0012 |
|  |  | | Postcentral_R | | | 36 -42 66 | | | -5.5 | | | 0.0012 |
|  |  | | Cingulum_Mid_R | | | 12 42 30 | | | -5.44 | | | 0.0012 |
|  |  | | Temporal_Pole_Sup_L | | | -54 6 -12 | | | -5.41 | | | 0.0012 |
|  |  | | Insula_L | | | -36 12 12 | | | -5.4 | | | 0.0012 |
|  |  | | SupraMarginal_R | | | 60 -24 24 | | | -5.4 | | | 0.0012 |
|  |  | | Occipital_Mid_L | | | -18 -96 0 | | | -5.39 | | | 0.0012 |
|  |  | | Frontal_Inf_Orb_R | | | 42 48 -12 | | | -5.38 | | | 0.0012 |
|  |  | | Occipital_Inf_L | | | -18 -96 -6 | | | -5.38 | | | 0.0012 |
|  |  | | Occipital_Sup_R | | | 18 -96 18 | | | -5.36 | | | 0.0012 |
|  |  | | Hippocampus_L | | | -24 -6 -24 | | | -5.29 | | | 0.0014 |
|  |  | | Parietal_Sup_R | | | 42 -48 60 | | | -5.29 | | | 0.0014 |
|  |  | | Lingual_L | | | -18 -96 -18 | | | -5.27 | | | 0.0014 |
|  |  | | Frontal_Mid_Orb_R | | | 42 54 -6 | | | -5.21 | | | 0.0014 |
|  |  | | Temporal_Sup_L | | | -54 6 -6 | | | -5.21 | | | 0.0014 |
|  |  | | Temporal_Pole_Sup_R | | | 42 18 -30 | | | -5.19 | | | 0.0014 |
|  |  | | Parietal_Inf_R | | | 42 -42 54 | | | -5.18 | | | 0.0014 |
|  |  | | Temporal_Pole_Mid_R | | | 48 12 -30 | | | -5.18 | | | 0.0014 |
|  |  | | Amygdala_L | | | -24 0 -24 | | | -5.17 | | | 0.0014 |
|  |  | | Cingulum_Ant_L | | | 0 36 30 | | | -5.15 | | | 0.0016 |
|  |  | | Cuneus_R | | | 18 -96 12 | | | -5.15 | | | 0.0016 |
|  |  | | Angular_L | | | -48 -72 24 | | | -5.15 | | | 0.0016 |
|  |  | | Occipital_Inf_R | | | 42 -66 -12 | | | -5.14 | | | 0.0016 |
|  |  | | Cingulum_Ant_R | | | 12 36 24 | | | -5.13 | | | 0.0016 |
|  |  | | Rectus_R | | | 6 30 -24 | | | -5.1 | | | 0.0016 |
|  |  | | Angular_R | | | 42 -60 54 | | | -5.08 | | | 0.0016 |
|  |  | | Rectus_L | | | 0 30 -24 | | | -5.06 | | | 0.0018 |
|  |  | | ParaHippocampal_R | | | 36 -30 -18 | | | -5.06 | | | 0.0018 |
|  |  | | Putamen_L | | | -24 12 12 | | | -5.01 | | | 0.0018 |
|  |  | | Amygdala_R | | | 36 0 -24 | | | -4.97 | | | 0.002 |
|  |  | | Cingulum_Mid_L | | | -6 24 36 | | | -4.95 | | | 0.0022 |
|  |  | | Frontal_Sup_Orb_R | | | 30 60 -6 | | | -4.94 | | | 0.0024 |
|  |  | | Frontal_Inf_Orb_L | | | -48 18 -6 | | | -4.91 | | | 0.0024 |
|  |  | | Caudate_L | | | -18 12 18 | | | -4.91 | | | 0.0024 |
|  |  | | Frontal_Sup_Orb_L | | | -12 12 -24 | | | -4.9 | | | 0.0024 |
|  |  | | Insula_R | | | 36 18 -18 | | | -4.88 | | | 0.0026 |
|  |  | | Olfactory_L | | | -24 6 -18 | | | -4.87 | | | 0.0026 |
|  |  | | Hippocampus_R | | | 36 -30 -12 | | | -4.85 | | | 0.0026 |
|  |  | | Lingual_R | | | 18 -48 -12 | | | -4.84 | | | 0.0026 |
|  |  | | Rolandic_Oper_R | | | 60 -18 18 | | | -4.82 | | | 0.002999 |
|  |  | | Occipital_Mid_R | | | 24 -96 6 | | | -4.77 | | | 0.003199 |
|  |  | | SupraMarginal_L | | | -60 -18 42 | | | -4.77 | | | 0.003199 |
|  |  | | Parietal_Inf_L | | | -54 -24 48 | | | -4.76 | | | 0.003199 |
|  |  | | Olfactory_R | | | 24 12 -18 | | | -4.74 | | | 0.003399 |
|  |  | | Frontal_Med_Orb_R | | | 12 54 -12 | | | -4.7 | | | 0.004199 |
|  |  | | Calcarine_R | | | 6 -90 6 | | | -4.7 | | | 0.003799 |
|  |  | | Caudate_R | | | 18 24 12 | | | -4.65 | | | 0.004599 |
|  |  | | Pallidum_L | | | -18 0 6 | | | -4.65 | | | 0.004599 |
|  |  | | Pallidum_R | | | 24 0 -6 | | | -4.65 | | | 0.004599 |
|  |  | | Putamen_R | | | 30 6 -6 | | | -4.64 | | | 0.004599 |
|  |  | | Thalamus_L | | | -12 -6 0 | | | -4.55 | | | 0.004799 |
|  | 0.25 Xenon | | Frontal_Inf_Orb_L | | | -36 48 -12 | | | -5.95 | | | 0.0007 |
|  |  | | Frontal_Mid_Orb_L | | | -36 54 -12 | | | -5.88 | | | 0.0008 |
|  |  | | Frontal_Sup_Orb_L | | | -24 48 -12 | | | -5.81 | | | 0.0008 |
|  |  | | Rectus_L | | | -12 48 -18 | | | -5.65 | | | 0.0016 |
|  |  | | Frontal_Inf_Tri_L | | | -48 42 12 | | | -5.6 | | | 0.002 |
|  |  | | Frontal_Mid_L | | | -36 48 12 | | | -5.56 | | | 0.0022 |
|  |  | | Paracentral_Lobule_L | | | -12 -24 78 | | | -5.56 | | | 0.0011 |
|  |  | | Frontal_Sup_Medial_L | | | -12 54 0 | | | -5.54 | | | 0.0022 |
|  |  | | Cingulum_Ant_L | | | -12 48 0 | | | -5.52 | | | 0.0022 |
|  |  | | Parietal_Inf_R | | | 48 -54 54 | | | -5.51 | | | 0.0022 |
|  |  | | Frontal_Sup_L | | | -18 54 12 | | | -5.47 | | | 0.0024 |
|  |  | | Frontal_Sup_Medial_R | | | 12 66 18 | | | -5.38 | | | 0.002799 |
|  |  | | Frontal_Sup_R | | | 18 66 12 | | | -5.32 | | | 0.002999 |
|  |  | | Rectus_R | | | 6 42 -24 | | | -5.31 | | | 0.003199 |
|  |  | | Frontal_Mid_R | | | 30 60 6 | | | -5.25 | | | 0.003199 |
|  |  | | Frontal_Med_Orb_R | | | 6 48 -12 | | | -5.24 | | | 0.003199 |
|  |  | | Cingulum_Ant_R | | | 6 42 0 | | | -5.17 | | | 0.003199 |
|  |  | | Frontal_Sup_Orb_R | | | 12 60 -18 | | | -5.13 | | | 0.003399 |
|  |  | | SupraMarginal_L | | | -60 -42 36 | | | -5.12 | | | 0.003399 |
|  |  | | Fusiform_L | | | -36 -18 -30 | | | -5.11 | | | 0.003599 |
|  |  | | Olfactory_L | | | -6 24 -12 | | | -5.1 | | | 0.003599 |
|  |  | | Temporal_Inf_L | | | -42 -24 -30 | | | -5.06 | | | 0.003799 |
|  |  | | Parietal_Inf_L | | | -60 -48 36 | | | -5.04 | | | 0.003799 |
|  |  | | Caudate_R | | | 6 18 0 | | | -5.02 | | | 0.003799 |
|  |  | | Temporal_Mid_L | | | -48 -18 -18 | | | -5.01 | | | 0.003799 |
|  |  | | Thalamus_L | | | -12 -6 6 | | | -4.92 | | | 0.003799 |
|  |  | | Insula_R | | | 30 -18 18 | | | -4.91 | | | 0.003999 |
