## Supplementary Digital Content 3 for "In search of Universal Cortical Power Changes Linked to NMDA-Antagonist based Anesthetic Induced Reductions in Consciousness"

Significant maximum statistics (p=0.025) corrected t-statistic maps of the power changes of N_2_O relative to Xe across subjects that demonstrate trends in the data at increasing equivalent gas concentrations of 0.25, 0.50, 0.75 MAC-awake in magnetoencephalography and electroencephalography datasets are shown in Supp. Figure 3.

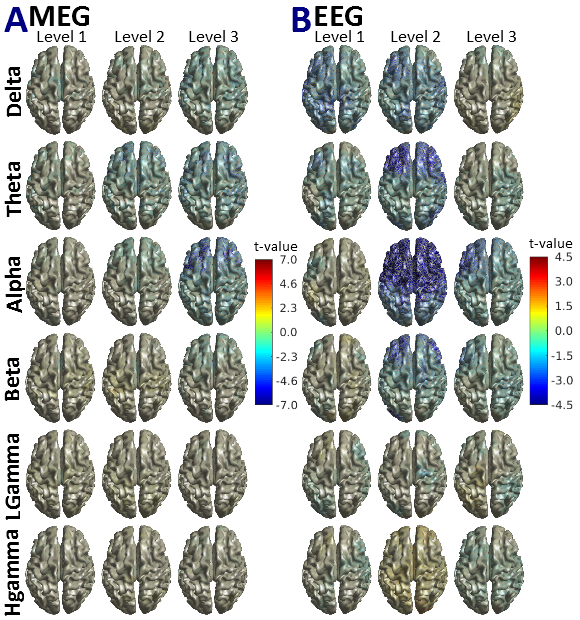

*Supp. Figure 3. Group level source power t-statistic maps contrasting equivalent doses of Xe and N_2_O.* The t-values for magnetoencephalographic (MEG - A) and electroencephalographic (EEG - B) point to subtle yet significant (p=0.05) changes in low frequency delta, theta and alpha when comparing the 0.25 (Level 1), 0.50 (Level 2) and 0.75 (Level 3) equi MAC-awake concentrations of Xe and N_2_O administered (N_2_O relative to Xe comparison). No significant differences appear across the two gases in high frequency beta and gamma activity. The difference in scale between A and B should be noted. [delta (1-4 Hz), theta (4-8 Hz), alpha (8-13 Hz), beta (13-30 Hz), Lgamma: low gamma (30-49 Hz), Hgamma: high gamma (51-99 Hz)].

Highly significant (p=0.004) power changes across the two gases in increasing gas levels contrasted to the post-antiemetic baseline reveal region specific changes in each frequency band investigate. Supp. Table 3 gives a full account of all significantly altered regions for N_2_O relative to Xe.

*Supp. Table 3. Magnetoencephalographic and Electroencephalographic sources most significantly altered in equivalent gas concentrations of Xe and N_2_O*. Significantly (p=0.004) changed regions of interest by contrasting equivalent inhaled concentrations of the two gases in magnetoencephalography and electroencephalography data. Voxel coordinates are in AAL atlas coordinate system along with associated labels^43^. [delta (1-4 Hz), theta (4-8 Hz), alpha (8-13 Hz), beta (13-30 Hz), Lgamma: low gamma (30-49 Hz), Hgamma: high gamma (51-99 Hz)].

| **Measurement** | **Frequency Band** | **MAC-awake Level** | | **Region of Interest** | **Voxel Coordinate** | **t-value** | **p-value** |
| --- | --- | --- | --- | --- | --- | --- | --- |
| **MEG** | **alpha** | | 0.75 | Frontal_Mid_R | 24 30 30 | -6.97 | 0.0017 |
|  |  | | 0.75 | Frontal_Inf_Oper_L | -48 6 18 | -6.86 | 0.0020 |
|  |  | | 0.75 | Frontal_Inf_Tri_L | -36 42 0 | -6.64 | 0.0032 |
|  |  | | 0.75 | Frontal_Mid_L | -24 30 30 | -6.42 | 0.0038 |
| **EEG** | **alpha** | | 0.50 | Frontal_Mid_R | 42 -6 54 | -6.22 | 0.0013 |
|  |  | | 0.50 | Frontal_Sup_Medial_L | -6 42 54 | -6.15 | 0.0016 |
|  |  | | 0.50 | Frontal_Mid_L | -24 30 54 | -6.14 | 0.0016 |
|  |  | | 0.50 | Supp_Motor_Area_L | -6 6 72 | -6.11 | 0.0018 |
|  |  | | 0.50 | Precentral_R | 42 -12 60 | -6.07 | 0.0022 |
|  |  | | 0.50 | Precentral_L | -24 -12 72 | -6.06 | 0.0024 |
|  |  | | 0.50 | Frontal_Sup_L | -18 42 48 | -5.98 | 0.0026 |
|  |  | | 0.50 | Cingulum_Mid_L | -6 12 42 | -5.84 | 0.0026 |
|  |  | | 0.50 | Frontal_Inf_Oper_L | -48 12 18 | -5.80 | 0.0026 |
|  |  | | 0.50 | Cingulum_Ant_L | -6 18 30 | -5.72 | 0.0028 |
|  |  | | 0.50 | Cingulum_Mid_R | 6 18 42 | -5.71 | 0.0028 |
|  |  | | 0.50 | Frontal_Sup_Medial_R | 6 24 42 | -5.69 | 0.0028 |
|  |  | | 0.50 | Supp_Motor_Area_R | 6 18 48 | -5.64 | 0.0028 |
|  |  | | 0.50 | Frontal_Sup_R | 18 30 60 | -5.63 | 0.0030 |
|  |  | | 0.50 | Frontal_Inf_Tri_L | -48 18 24 | -5.62 | 0.0030 |
|  |  | | 0.50 | Insula_L | -36 12 12 | -5.56 | 0.0030 |
|  |  | | 0.50 | Cingulum_Ant_R | 6 18 24 | -5.27 | 0.0056 |
|  |  | | 0.50 | Caudate_L | -18 6 24 | -5.21 | 0.0058 |
|  |  | | 0.50 | Rolandic_Oper_L | -42 0 18 | -5.18 | 0.0058 |
|  |  | | 0.50 | Frontal_Inf_Orb_L | -36 24 -6 | -5.14 | 0.0058 |
|  |  | | 0.50 | Paracentral_Lobule_L | -18 -12 66 | -5.00 | 0.0066 |
|  |  | | 0.50 | Putamen_L | -24 12 12 | -4.96 | 0.0072 |
|  |  | | 0.50 | Temporal_Pole_Mid_R | 42 18 -36 | -4.96 | 0.0072 |
|  |  | | 0.50 | Frontal_Inf_Oper_R | 30 12 30 | -4.95 | 0.0072 |
|  |  | | 0.50 | Temporal_Pole_Sup_R | 42 18 -30 | -4.89 | 0.0080 |
|  |  | | 0.50 | Postcentral_L | -60 -6 30 | -4.88 | 0.0082 |
|  |  | | 0.50 | Caudate_R | 18 6 24 | -4.88 | 0.0080 |
|  |  | | 0.50 | Postcentral_R | 48 -6 30 | -4.83 | 0.0084 |
|  |  | | 0.50 | Temporal_Pole_Sup_L | -30 18 -30 | -4.83 | 0.0084 |
|  |  | | 0.50 | Insula_R | 42 18 -12 | -4.81 | 0.0084 |
|  |  | | 0.50 | Frontal_Inf_Tri_R | 36 24 12 | -4.78 | 0.0086 |
|  |  | | 0.50 | Frontal_Inf_Orb_R | 42 24 -18 | -4.78 | 0.0086 |
|  |  | | 0.50 | Temporal_Pole_Mid_L | -36 18 -36 | -4.71 | 0.0088 |
|  |  | | 0.50 | Temporal_Sup_R | 60 0 -6 | -4.66 | 0.0092 |
|  |  | | 0.50 | Temporal_Inf_L | -42 6 -42 | -4.65 | 0.0094 |
|  |  | | 0.50 | Rolandic_Oper_R | 42 6 12 | -4.64 | 0.0094 |
|  |  | | 0.50 | Putamen_R | 30 18 0 | -4.63 | 0.0094 |
|  |  | | 0.50 | Frontal_Mid_Orb_L | -24 30 -18 | -4.62 | 0.0100 |
|  |  | | 0.50 | Olfactory_L | -18 12 -18 | -4.62 | 0.0098 |
|  |  | | 0.50 | ParaHippocampal_R | 24 12 -30 | -4.58 | 0.0108 |
|  |  | | 0.50 | Temporal_Inf_R | 36 6 -42 | -4.48 | 0.0122 |
|  |  | | 0.50 | Rectus_L | -12 18 -12 | -4.47 | 0.0122 |
|  |  | | 0.50 | Fusiform_R | 24 6 -42 | -4.47 | 0.0122 |
|  |  | | 0.50 | Fusiform_L | -30 0 -36 | -4.44 | 0.0132 |
|  |  | | 0.50 | ParaHippocampal_L | -18 6 -24 | -4.42 | 0.0142 |
|  |  | | 0.50 | Temporal_Mid_L | -36 6 -30 | -4.42 | 0.0140 |
|  |  | | 0.50 | Frontal_Sup_Orb_L | -12 18 -18 | -4.41 | 0.0142 |
|  |  | | 0.50 | Amygdala_L | -24 0 -12 | -4.40 | 0.0142 |
|  |  | | 0.50 | Olfactory_R | 24 12 -18 | -4.39 | 0.0148 |
|  |  | | 0.50 | Temporal_Mid_R | 48 6 -30 | -4.39 | 0.0148 |
|  |  | | 0.50 | Heschl_R | 42 -24 12 | -4.38 | 0.0150 |
|  |  | | 0.50 | Amygdala_R | 24 6 -18 | -4.36 | 0.0154 |
|  |  | | 0.50 | Pallidum_L | -18 6 0 | -4.36 | 0.0152 |
|  |  | | 0.50 | Frontal_Mid_Orb_R | 42 48 -6 | -4.35 | 0.0156 |
|  |  | | 0.50 | Temporal_Sup_L | -48 6 -6 | -4.31 | 0.0158 |
|  |  | | 0.50 | Pallidum_R | 24 0 0 | -4.27 | 0.0164 |
|  |  | | 0.50 | Frontal_Sup_Orb_R | 24 30 -24 | -4.23 | 0.0178 |
|  |  | | 0.50 | Thalamus_R | 12 -12 18 | -4.21 | 0.0178 |
|  |  | | 0.50 | SupraMarginal_R | 60 -42 24 | -4.14 | 0.0188 |
|  |  | | 0.50 | Paracentral_Lobule_R | 6 -42 78 | -4.14 | 0.0188 |
|  |  | | 0.50 | Rectus_R | 18 18 -18 | -4.13 | 0.0190 |
|  |  | | 0.50 | Hippocampus_R | 18 6 30 | -4.10 | 0.0192 |
|  |  | | 0.50 | Thalamus_L | -6 -12 18 | -4.05 | 0.0198 |
|  |  | | 0.50 | Hippocampus_L | -18 -6 -12 | -4.03 | 0.0204 |
|  |  | | 0.50 | Parietal_Sup_R | 18 -48 72 | -4.03 | 0.0206 |
|  |  | | 0.50 | Parietal_Sup_L | -24 -54 72 | -4.01 | 0.0206 |
|  |  | | 0.50 | Frontal_Med_Orb_L | -12 36 -12 | -3.93 | 0.0226 |
|  |  | | 0.75 | Frontal_Mid_L | -42 48 18 | -3.71 | 0.0141 |
|  |  | | 0.75 | Frontal_Inf_Tri_L | -54 24 12 | -3.45 | 0.0230 |
|  | **theta** | | 0.50 | Frontal_Inf_Tri_L | -48 36 24 | -4.66 | 0.0111 |
|  |  | | 0.50 | Frontal_Inf_Tri_R | 54 30 24 | -4.37 | 0.0156 |
|  |  | | 0.50 | Frontal_Mid_L | -48 30 30 | -4.36 | 0.0156 |
|  |  | | 0.50 | Frontal_Mid_R | 36 24 54 | -4.11 | 0.0208 |
|  | **beta** | | 0.50 | Frontal_Mid_R | 42 48 24 | -4.61 | 0.0139 |
|  |  | | 0.50 | Frontal_Inf_Tri_R | 48 36 24 | -4.41 | 0.0184 |
|  |  | | 0.50 | Occipital_Mid_L | -36 -90 6 | -4.37 | 0.0192 |
|  | **delta** | | 0.50 | Precentral_R | 60 -6 42 | -3.95 | 0.0231 |
